# A metabolomics pipeline enables mechanistic interrogation of the gut microbiome

**DOI:** 10.1101/2021.05.25.445684

**Authors:** Shuo Han, Will Van Treuren, Curt R. Fischer, Bryan D. Merrill, Brian C. DeFelice, Juan M. Sanchez, Steven K. Higginbottom, Leah Guthrie, Lalla A. Fall, Dylan Dodd, Michael A. Fischbach, Justin L. Sonnenburg

## Abstract

Gut microbes modulate host phenotypes and are associated with numerous health effects in humans, ranging from cancer immunotherapy response to metabolic disease and obesity. However, difficulty in accurate and high-throughput functional analysis of human gut microbes has hindered defining mechanistic connections between individual microbial strains and host phenotypes. One key way the gut microbiome influences host physiology is through the production of small molecules^1–3^, yet progress in elucidating this chemical interplay has been hindered by limited tools calibrated to detect products of anaerobic biochemistry in the gut. Here we construct a microbiome-focused, integrated mass-spectrometry pipeline to accelerate the identification of microbiota-dependent metabolites (MDMs) in diverse sample types. We report the metabolic profiles of 178 gut microbe strains using our library of 833 metabolites. Leveraging this metabolomics resource we establish deviations in the relationships between phylogeny and metabolism, use machine learning to discover novel metabolism in *Bacteroides*, and employ comparative genomics-based discovery of candidate biochemical pathways. MDMs can be detected in diverse biofluids in gnotobiotic and conventional mice and traced back to corresponding metabolomic profiles of cultured bacteria. Collectively, our microbiome-focused metabolomics pipeline and interactive metabolomics profile explorer are a powerful tool for characterizing microbe and microbe-host interactions.

## Introduction

The human gut microbiota encode diverse metabolic pathways. Enriched in anaerobic pathways that process diverse diet- and host-derived molecules, gut microbes make numerous novel compounds with relevance for human health and untapped therapeutic potential. Many of these microbial products in the gut subsequently enter the host’s tissue and circulation, where additional metabolic steps can add to the chemical diversity^1–3^. Several recent studies have shown that microbiota-dependent metabolites (MDMs) influence immune function^4^, metabolism^5, 6^, cardiovascular health^7^, and cognition and behavior^8^. In many cases, MDMs exert these effects on host biology by binding to specific host receptors^9^ and activating downstream signaling pathways^10^. Discovery of how individual prevalent human gut microbes mechanistically contribute to host phenotypes has been hampered by the difficulty in accurately monitoring the diverse universe of molecules produced by gut microbes. To address this gap, recent studies in the field have leveraged improvements in high resolution mass spectrometry^11^ as well as growing mass spectra and compound databases^12^ (e.g. MoNA, METLIN^13^, HMDB^14^, and KEGG^15^). Nevertheless, because of 1) fundamental differences between anaerobic metabolism in the gut vs. aerobic biochemistry, and 2) under-representation of anaerobic microbial products in existing databases, the full metabolic capability of the microbiota remains understudied. Here we present a microbiome-focused, integrated mass-spectrometry pipeline to facilitate the identification of microbiota-dependent metabolites in diverse sample types, and to associate these metabolites with microbial strains and genetic pathways.

### Microbiome-focused metabolomics

To enable interrogation of microbiome metabolism, we 1) constructed a mass spectrometry-based reference library to detect anaerobic biochemistry and an analytical pipeline to integrate large metabolomic datasets; 2) validated our methods to ensure applicability to the broader scientific community; and 3) enabled interactive, public access to our datasets (https://sonnenburglab.github.io/Metabolomics_Data_Explorer) (Fig. 1, Extended Data Fig. 1-3, Supplementary Table 1-4, Methods).

**Fig. 1.**
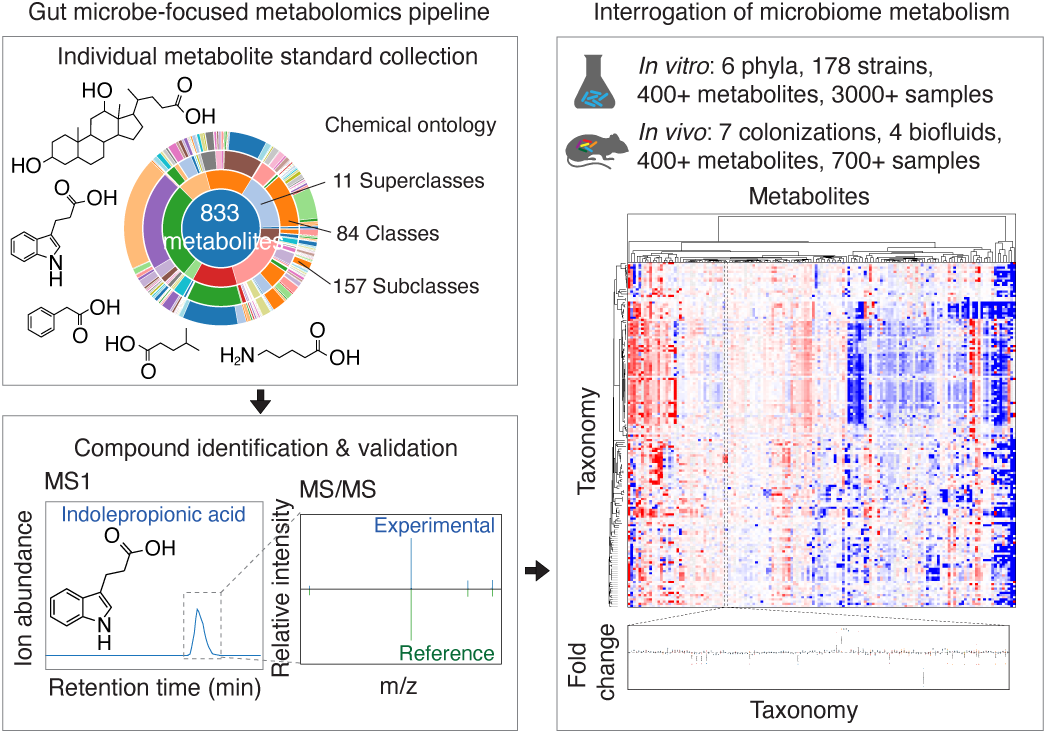
A microbiome-focused metabolomics pipeline enables mechanistic interrogation of microbiome metabolism. Schematic of our metabolomics workflow, consisting of MS reference library construction and validation, producing *in vitro* and *in vivo* metabolomic profiles across diverse sample types. Our entire dataset is publicly accessible via a web-based, interactive Metabolomics Data Explorer.

Next, we leveraged this tool to create a reference dataset of metabolomic profiles for individual bacterial strains to enable multiple modes of analysis and discovery. 178 individual prevalent human gut microbes representing 130 species and spanning 6 phyla were acquired from ATCC, DSMZ, and BEI (Supplementary Table 5, 6). To create the most comparable dataset of metabolism, we cultivated all supported strains (158/178) in mega medium – a rich, undefined medium known to support growth of diverse bacteria and collected culture supernatant between mid-log to stationary phase (Extended Data Fig. 4a, b, Supplementary Methods). The remaining 20 strains were grown in 9 additional media as detailed (Supplementary Table 6), 29 strains were grown and analyzed across multiple media types (Supplementary Table 7, Extended Data Fig. 4c).

To assess large-scale metabolite production and consumption patterns, we hierarchically clustered individual bacterial strains (Extended Data Fig. 4d-f, Supplementary Table 7). In some cases, two closely related species exhibiting distinct metabolomic profiles punctuated with metabolite-level similarities (*Clostridium sporogenes* and *Clostridium cadaveris,* Extended Data Fig. 5a, b). In others, phylogenetic proximity is accompanied by similarity in metabolic patterns (four *Bacteroides fragilis* strains, Pearson r > 0.80 for all pairwise comparisons, Extended Data Fig. 5a, b). Conversely, hierarchical clustering of species by metabolomic profile distance reveals unexpectedly shared metabolic patterns among phylogenetically distant species (*Atopobium parvulum,* Phylum Actinobacteria and *Catenibacterium mitsuokai*, Phylum Firmicutes) (Extended Data Fig. 6a-c).

In addition to the large-scale metabolic patterns, we discovered unique high producers or consumers of specific metabolites within our strain collection. For example, *Enterococcus faecalis* and *Enterococcus faecium* significantly produce high levels of tyramine (Extended Data Fig. 4e), a biogenic amine known to modulate host neurological functions^16^. In contrast, *Clostridium cadaveris* significantly consumes high levels of pantothenic acid (vitamin B5) (Extended Data Fig. 4f), a molecule associated with inflammatory bowel diseases^17^. This large-scale *in vitro* screen enables us to identify numerous high-abundance, variably conserved, microbially-derived metabolites that can be tracked *in vitro* and *in vivo* (Extended Data Fig. 6d).

### Relationships between phylogeny, taxonomy, and metabolome

We next addressed large-scale relationships between strain metabolism and phylogeny, a complex topic addressed with different approaches in previous studies^18–21^. Bacterial metabolism is a product of a microbe’s genetic metabolic toolkit and the chemical environment. Comparing metabolomic and phylogenetic trees for the same set of 158 mega-medium grown strains revealed broadly conserved topology with the strains most often clustering by phyla (Fig. 2a, Extended Data Fig. 6a, 7a, Supplementary Methods). However, this similarity is punctuated by significant divergences where the relative location of specific strains in the two trees differs substantially (magenta and gold colored branches, Fig. 2a). Notably, these patterns of clustering are preserved when metabolites are weighted by chemical similarity (Extended Data Fig. 7b, c; Mantel test: r^2^ = 0.863, *P* = 0.001).

**Fig. 2.**
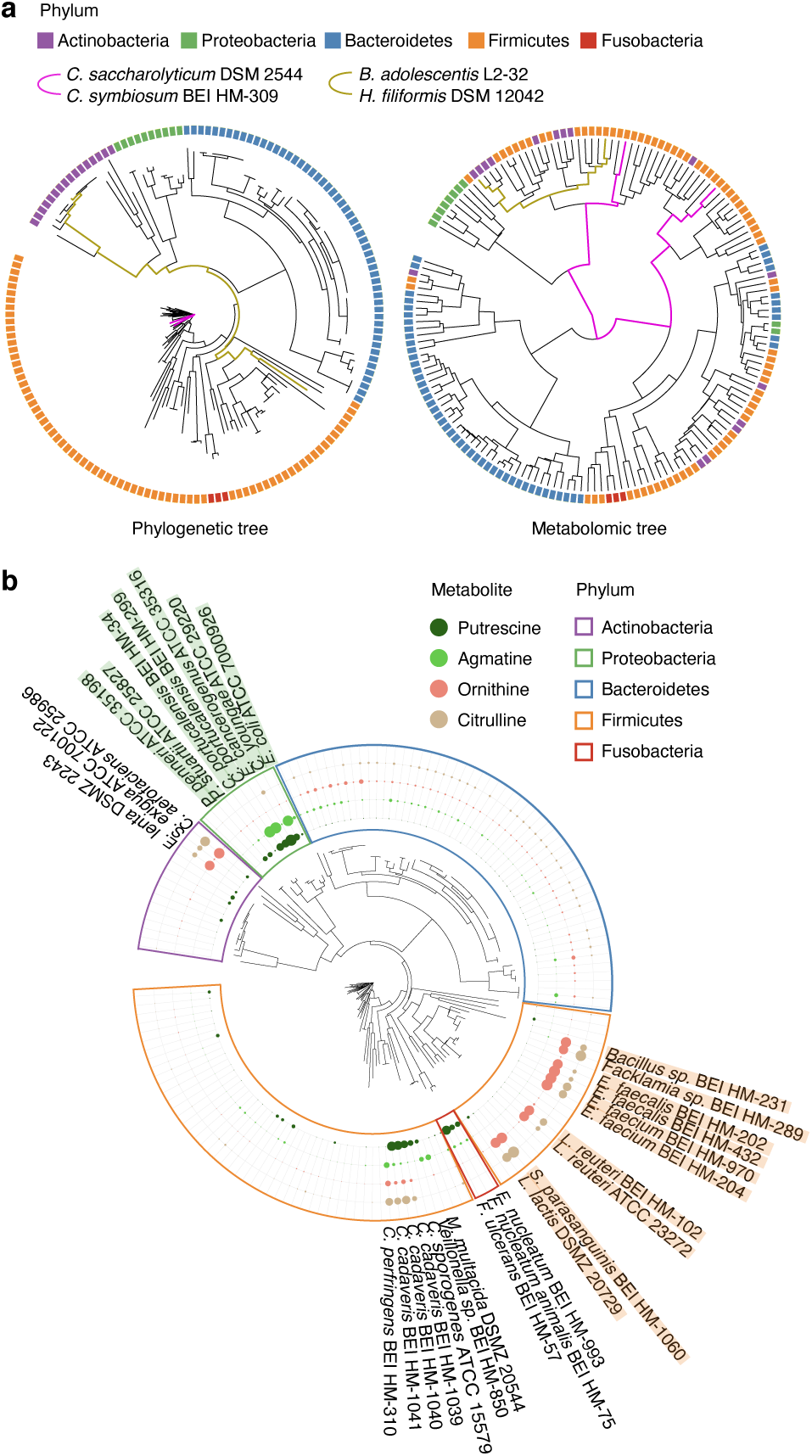
Relationships between phylogeny, taxonomy, and metabolome. **a,** Comparison of tree topology constructed based on phylogenetic (left) and metabolomic profile (fold-change data, right) distance matrices of 158 mega-medium grown strains spanning five phyla (one to three independent experiments, each with n ≥ 3 biological replicates). **b,** Metabolite accumulation patterns across all 158 mega-medium grown strains, clustered based on phylogenetic distance. Dot size: mean production levels of one to three independent experiments, each with n ≥ 3 biological replicates. For each metabolite, the largest dot represents the highest production level for that metabolite.

To quantify these differences we compared metabolomic distance between strains to their evolutionary distance (Extended Data Fig. 7d, Supplementary Table 7). Using V4 16S the relationship between phylogenetic distance and metabolomic distance is linear (r^2^ = 0.30, *P* < 1e- 92) below ∼0.11 branchlength units, approximating a difference of taxonomic ’Class’ in our data. Above a branchlength of 0.11, 16S distance explains almost none of the variance in metabolomic distance (r^2^ = 0.02, *P* < 10^-9^). These patterns are robust to data transformation and evolutionary distance derived from full-length 16S genes (Extended Data Fig. 7e-j). Comparing metabolic distance of bacteria grouped by taxonomic rank alone (e.g., distance between different strains of the same species) reveals a similar pattern of saturation (Extended Data Fig. 7d, Supplementary Table 7). These data indicate that when two strains are grown in the same complex medium, differences in detected microbial metabolism are smaller on average than what would be extrapolated linearly from evolutionary or taxonomic relationships, particularly for distantly related bacteria. Importantly, the high variance in metabolic distance between microbes of any relatedness (taxonomic or phylogenetic) reaffirms the utility of metabolite profiles when comparing specific strains.

We next leveraged our strain-resolved metabolomic and genomic data to examine the correlation between bacterial genetic and metabolic variations in the context of a single pathway, polyamine biosynthesis (Fig. 2b, Extended Data Fig. 7k). Gut microbially-derived putrescine and its precursor ornithine have both been implicated in influencing aspects of host physiology^22, 23^. Their biosynthetic enzymes have been functionally characterized in select bacterial species (e.g., ornithine-producing *arc* genes^24^, putrescine-producing *spe* genes^25^).

We discovered two groups of phylogenetically distant strains in two phyla, Firmicutes and Actinobacteria (Fig. 2b, orange and purple phylum borders respectively), that accumulate high levels of ornithine and citrulline in the absence of significant downstream polyamine production. We performed iterative comparative genomics starting with the ornithine-producing *arc* genes described in *Enterococcus faecalis* and found their conserved presence (Extended Data Fig. 7k) among the ornithine-accumulating strains, such as the Lactobacillales (Fig. 2b, strain names highlighted in orange). Notably, these genes are not detectable in the non-ornithine-accumulating phylogenetic neighbors, for both the Lactobacillales and Actinobacteria. These examples illustrate that when metabolic phenotypes depart from phylogeny, orthologous gene-to-metabolite relationships may be preserved. We next identified strains that accumulate high levels of downstream putrescine and/or agmatine within three phyla: Proteobacteria, Fusobacteria, and Firmicutes (Fig. 2b, green, red, and orange phylum borders respectively). While several putrescine-accumulating Proteobacteria strains (Fig. 2b, strain names highlighted in green) share the putrescine-producing *spe* gene cluster described in *E. coli* (Extended Data Fig. 7k), these genes are not detectable in the Fusobacteria. These data indicate limited ability of phylogeny or genome-based prediction of metabolic functions in bacterial strains and reinforce the utility of measuring metabolic phenotypes to identify strains and genes producing specific metabolites that have the potential to impact host biology.

### Identifying novel gene-phenotype relationships

Metabolite production and consumption have long been used as mechanisms to group and identify organisms (e.g., indole production). Here, we used our comprehensive mega medium-derived metabolomic data along with simple machine learning (random forest; RF) models to identify sets of metabolites distinguishing different taxonomic groups. Simple RF models could accurately classify the taxonomic origin of microbial supernatants (Fig. 3a, Supplementary Methods). While the total metabolome is not clearly predictive of taxonomy (Fig. 2a, Extended Data Fig. 7d) these RF’s revealed subsets of the chemical features that were highly conserved and predictive of taxonomic identity (Extended Data Fig. 8a).

**Fig. 3.**
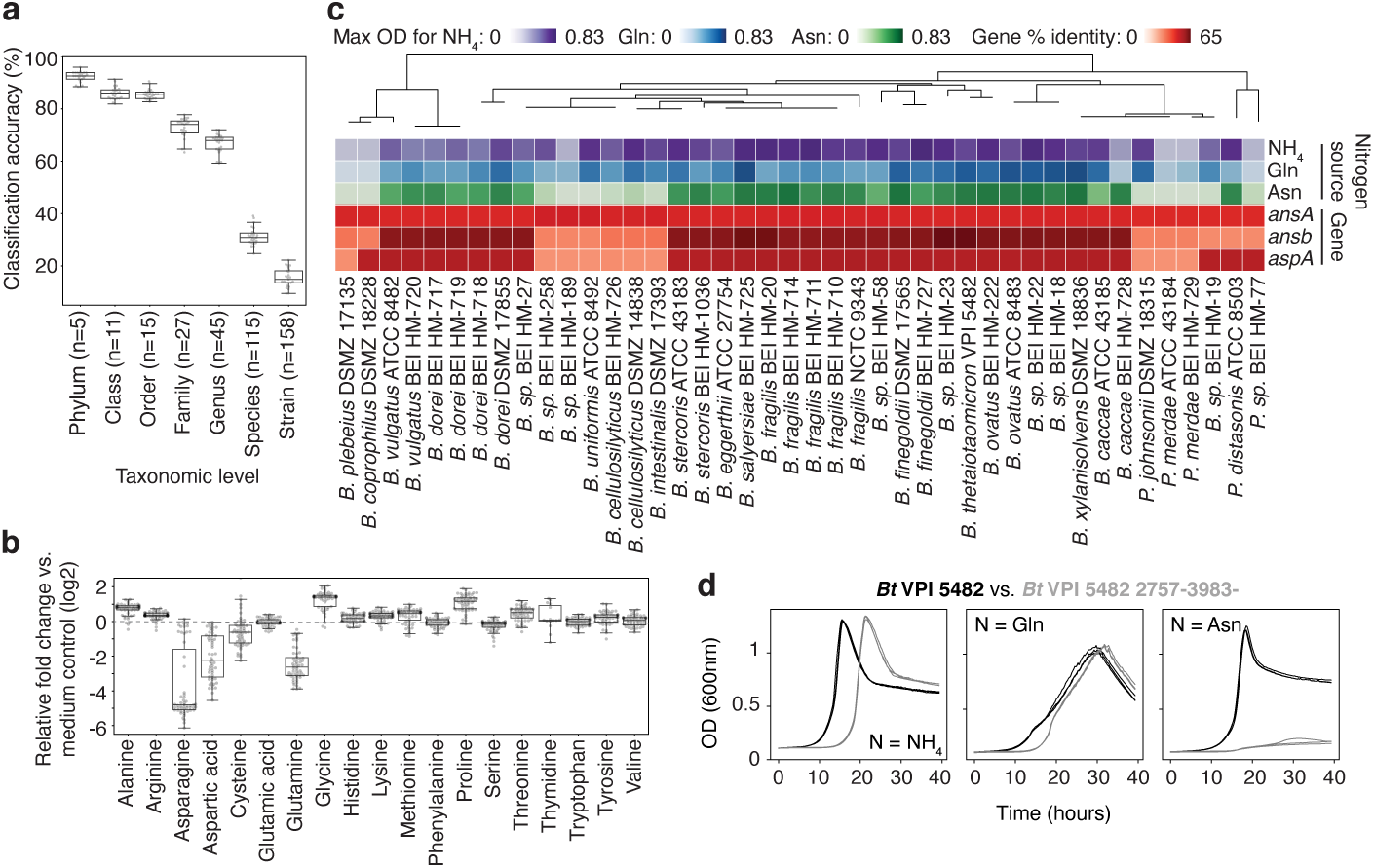
Discovery of nitrogen assimilation strategies in Bacteroides and novel gene-phenotype relationships. **a,** Classification accuracy of Random Forest (RF) models at each taxonomic level, based on metabolomic profiles of 158 mega-medium grown bacterial strains from one to three independent experiments, each with n ≥ 3 biological replicates. Phylum (n = 5), Class (n = 11), Order (n = 15), Family (n = 27), Genus (n = 45), Species (n = 115), and Strain (n = 158). **b,** Amino acid production or consumption levels by Bacteroidetes strains from one to three independent experiments, each with n ≥ 3 biological replicates. Only uniquely detected (non-coeluting) amino acids are shown. **a**, **b**, Boxes: median, 25th, and 75th percentile; whiskers: Tukey’s method. **c,** Phylogenetic tree of Bacteroidetes strains, growth curve max optical density (OD_600nm_), and percentage of protein sequence identity for *E. coli* asparagine-consuming, ammonium-liberating enzymes. Nitrogen sources: ammonia (NH_4_), Glutamine (Gln), Asparagine (Asn). **d,** Representative growth curves of wild-type and mutant *Bt* (2757-3983-) in modified Salyer’s Minimal Medium from one experiment with n = 3 biological replicates. Nitrogen sources as in **c**: NH4, Gln, Asn.

The most discriminating features selected by the RFs for differentiating phyla had an overrepresentation in amino acid (AA) metabolism (Extended Data Fig. 8a). Intriguingly, Bacteroidetes were differentiated by their consumption of most of the glutamine (Gln; median consumption 83%) and asparagine (Asn; median consumption 96%) in the mega medium (Fig. 3b). Classic work showing Bacteroides could not use free AAs as sole nitrogen sources (NS) failed to test Asn and Gln^26^. Based on the data from the 60 Bacteroidetes taxa in the collection, we hypothesized that Gln and Asn might serve as sole NS. To test this, we grew all 60 Bacteroides and Parabacteroides species in a minimal medium that lacked free ammonium, but contained 10 mM Glutamate (Glu), Gln, or Asn. Strikingly, the Asn or Gln sufficed as the NS for 50 of 60 Bacteroidetes tested (Fig. 3c, Extended Data Fig. 8b, c). To determine the genetic basis of Asn utilization, we searched the Bacteroidetes genomes for homologs of *E. coli* enzymes that consume asparagine and release ammonia (Fig. 3c, red rows). For taxa with available genomes, an L-asparaginase II homolog (*ansB*; > 59% identity) strongly correlated (Pearson r = 0.91) with max OD when grown on Asn. Using a transposon mutant in the *Bacteroides thetaiotaomicron* type strain (*Bt* VPI 5482 2757-3983-), we confirmed that this L-asparaginase II homolog was necessary for growth with Asn as a sole NS (Fig. 3d). The effect we observed was not dependent on presence of cysteine; *Bt* VPI 5482 and *Bt* VPI 5482 2757-3983- grew with sodium sulfide substituted as reduced sulfur source and the pattern of growth was maintained (Extended Data Fig. 8d). We next examined Bacteroides amino acid consumption patterns *in vivo*. In the cecum of mice monocolonized with *Bt* VPI 5482, Asn was the most depleted AA (median decrease of 86.9%) compared to germ-free control animals (Extended Data Fig. 8e). This observation is consistent with *in vivo* Asn utilization by *Bt*, but does not exclude colonization-dependent changes in host Asn utilization. These findings demonstrate the power of combining strain-resolved metabolomics with simple statistical models, in this case to discover a major metabolic capacity for nitrogen assimilation for the most abundant genus in the industrialized microbiota.

### Individual microbial contribution in a multi-species community

Pursuit of mechanism in microbiome studies can be aided by reverse translation of findings from complex communities (humans or conventional animals) into highly controlled (e.g., gnotobiotic) models. We have recently demonstrated the utility of our *in vitro* strain metabolite profiles in reverse translation by recreating metabolic phenotypes of interest to study IBD mechanisms^27^. Based on two metabolites detected in human biofluids^28^ and conventional mice, we asked whether we could reconstitute the production of microbially-derived metabolites in the host gut and/or circulation by colonizing mice with the highest *in vitro* producing strain in our collection. One candidate, agmatine, is a polyamine with neuroprotective roles in mammals^29^, and a substrate for transporters in kidney and liver cells^30^. The other candidate, alpha-ketoglutaric acid, is a tricarboxylic acid cycle intermediate that extends lifespan in nematode *C. elegans* and increases autophagy in mammalian cells^31^.

Consistent with our *in vitro* observations, agmatine and alpha-ketoglutaric acid levels were both significantly increased in the feces of mice mono-associated with a high *in vitro* producer: *Citrobacter portucalensis* and *Anaerostipes sp.*, respectively (Fig. 4a, Extended Data Fig. 9a). Further, we bolstered agmatine levels in the host circulation (e.g., urine) relative to the germ-free controls (Fig. 4a). These examples provide a proof-of-concept application of our *in vitro* dataset to reconstitute specific microbially-derived metabolism in a mouse model, enabling potential mechanistic studies relevant to host physiology.

**Fig. 4.**
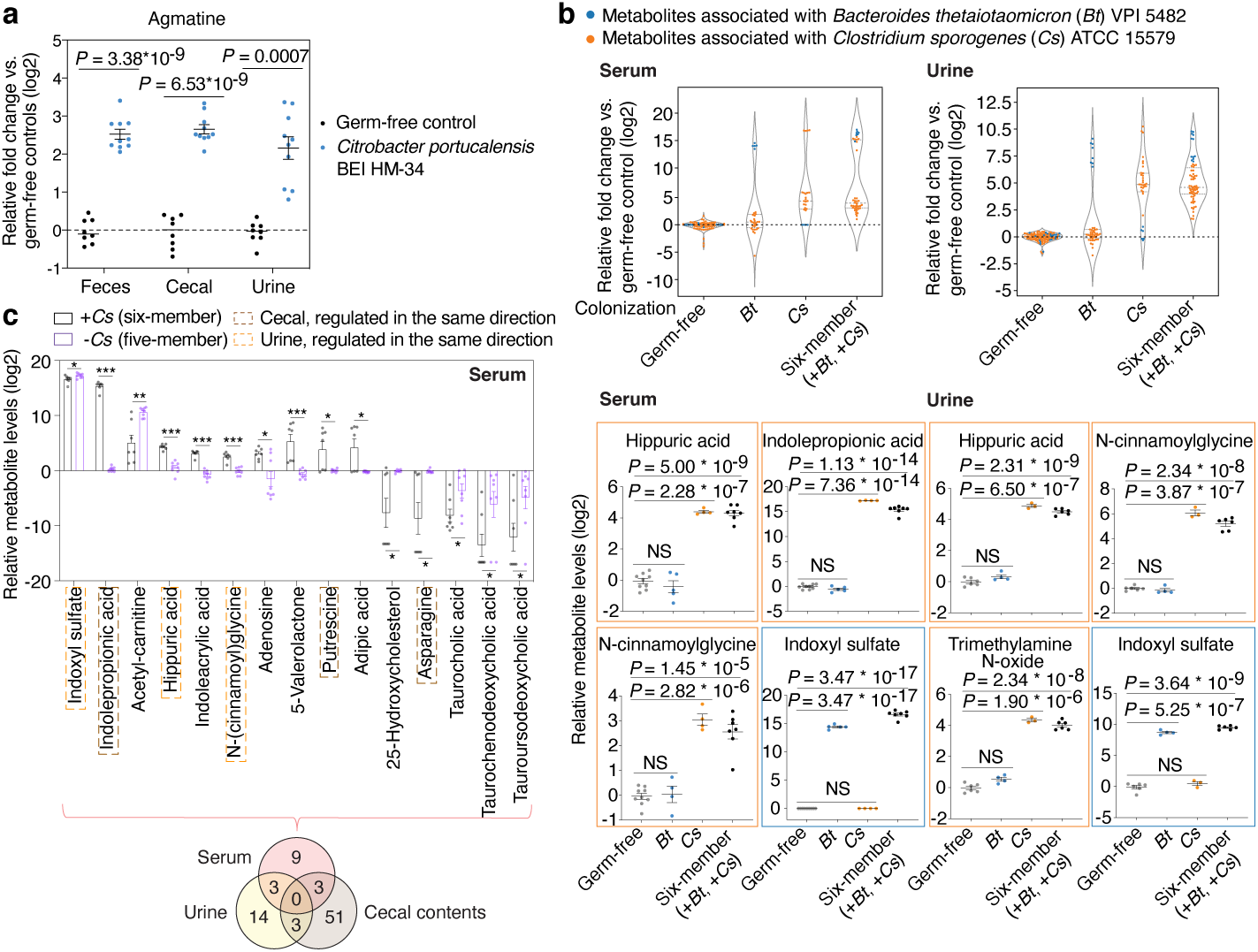
Metabolic contribution by individual gut microbes in a multi-species community. **a,** Quantification of agmatine levels. Mean ± s.e.m. of two independent experiments, each with n = 4 (germ-free) or n = 5 (*Citrobacter* mono-colonized) individual mice. **b**, Significantly produced metabolites associated with *Clostridium sporogenes* (*Cs*) or *Bacteroides thetaiotaomicron* (*Bt*) in serum (left panel) or urine (right panel). Violin plot: median and quartiles. Mean ± s.e.m. of one experiment with n = 4 (*Cs*, serum), n = 3 (*Cs*, urine), n = 5 (*Bt*, serum), n = 4 (*Bt*, urine), n = 7 (six-member, serum), n = 6 (six-member, urine), and n =9 germ-free mice pooled from both mono-association (n = 4) and community (n = 5) experiments. **c**, Serum metabolite levels in mice colonized with the six-member community (+*Cs*) or the five-member community (-*Cs*). Metabolites shown represent a panel of significantly elevated or reduced metabolites (≥ 4-fold, corrected *P* < 0.05) in the six-member community. Mean ± s.e.m. of one experiment with n = 7 (six-member community) and n = 8 (five-member community) mice. Venn diagram: significantly elevated or reduced metabolites in different host biofluids based on the same threshold defined above. **a**-**c**, *P* values: two-tailed *t*-test with Benjamini-Hochberg correction for multiple comparisons. * *P* < 0.05, ** *P* < 0.01, *** *P* < 0.001.

We leveraged our unique strain-resolved metabolomic data set combined with gnotobiotic colonization (Supplementary Table 8) and asked whether specific *in vivo* gut bacteria-derived metabolites serve as biomarkers for a given taxonomic group. Among the 34 significantly produced metabolites in both colonized mice and individual strain cultures, we found several phylum-specific metabolites (e.g., 5-Aminopentanoic acid and Indolepropionic acid by Firmicutes; Malic acid and Melatonin by Bacteroidetes) (Extended Data Fig. 9b, Supplementary Table 9). These data highlight that taxa-specific metabolites may serve as biomarkers for aspects of microbiome composition.

We next assessed the extent to which metabolites produced *in vitro* are reconstituted in gnotobiotic mice colonized with the same microbes. At the metabolomic profile level, *Clostridium sporogenes* (*Cs*)- or *Bacteroides thetaiotaomicron* (*Bt*)-mono-associated murine feces and cecal contents correlated with *Cs* or *Bt in vitro* culture when compared against 158 mega-medium grown taxa (*Cs*, top 1%; *Bt*, top 10%; Extended Data Fig. 9c). The lack of correlation in serum and urine (∼0, Extended Data Fig. 9c) is likely due to the inability of bacterial culture to recapitulate host-encoded metabolism (e.g., phase I/II enzymes). At the individual metabolite level, 8 out of 20 (40%, *Cs*) and 3 out of 29 (10%, *Bt*) significantly produced cecal metabolites *in vivo* were also produced by the same strain *in vitro* (Extended Data Fig. 9d). Further, when assessing a six-species defined microbiota, 15 out of 46 (33%) significantly produced cecal metabolites were also produced by one or more of the six species *in vitro* (Extended Data Fig. 9d). Collectively, these data illustrate that metabolites produced in a standard rich medium can inform a portion of the microbially-derived metabolites produced in the gut environment.

To better understand whether and how microbe-dependent metabolites in the gut can inform circulating metabolites in the host, we examined enteric and systemic metabolic contributions in the host by *Cs* and *Bt*. We measured metabolite profiles of four sample types (feces, cecal contents, serum, and urine) in different colonization states (Fig. 4b). Principal Component Analyses (PCA) reveal that metabolomic profiles cluster by sample types (e.g., cecal contents vs. serum) from mice colonized with the same microbes, as well as by colonization states (e.g., *Cs* mono-association vs. *Cs-*containing six-member community) (Extended Data Fig. 9e, f). We identified a distinct set of known and candidate host-microbial co-metabolites that are significantly elevated in the serum and/or urine, and are strongly associated with the presence of either *Cs* or *Bt* in the gut (Fig. 4b, Extended Data Fig. 9g, h). Notably, in both serum and urine, accumulation of N-(cinnamoyl)glycine is *Cs*-dependent, whereas accumulation of Indoxyl sulfate is *Bt*-dependent (Fig. 4b, Extended Data Fig. 9g, h). Our systematic and high-throughput detection of microbe and host-microbe metabolites across different sample types (e.g., from cecum to serum) enables identification of intermediates within known or candidate host-microbial co-metabolism pathways (Extended Data Fig. 10a).

To determine whether enteric presence of *Cs* is necessary for the elevation or reduction in specific metabolites in host circulation, we omitted *Cs* from the original six-member community. Metabolites shown are significantly elevated or reduced by at least four-fold in the serum, urine, or cecal contents of mice with the six-member community, relative to their germ-free controls (Fig. 4c, Extended Data Fig. 10b, c). In contrast, the five-member, *Cs*-lacking community either abrogated the production or restored the depletion of a subset of these metabolites in the serum or urine, indicating that enteric presence of *Cs* is necessary for modulating levels of these metabolites in the host circulation (Fig. 4c, Extended Data Fig. 10b) and illustrating the potential of microbiome editing to alter MDMs that circulate in the host blood.

## Discussion

Untargeted metabolomics has led to many discoveries of microbiota-dependent metabolic pathways^9, 10^ and metabolites linked to host diseases^17, 32–34^, yet there is exceptional untapped potential. Here we present a customizable and expandable method of constructing a chemical standard library-informed metabolomics pipeline tailored to detecting products of gut anaerobic biochemistry. Using this method, we construct an atlas of gut microbiota-dependent metabolic activities *in vitro* and *in vivo,* enabling functional studies of gut microbial communities. Complementary to recent studies using phylogenetic (16S)^35^ or metagenomic comparisons^36^ to predict gene functions, we used strain-resolved metabolomics to provide expansive biochemical profiles of individual strains. These profiles demonstrate that substantial metabolic variation is common even between closely related strains. Our findings, along with emerging studies on microbiome-focused metabolomics^37–39^ and gut microbial metabolism^40, 41^, reinforce the limits of phylogeny or genome-scale analysis to provide direct measurement or prediction of metabolic phenotypes and the molecules that link the microbiota to host physiology. Our existing strain-specific genome-by-metabolic profile data provides a rich resource for comparative discovery of genes and pathways that underlie bacterial phenotypic variation. Furthermore, this data and approach can be used as a direct reference or readily implemented platform for improving MDM identification in biological samples. Adding novel microbially-derived metabolites, along with new strains such as those isolated from diverse human populations, will uncover new mediators of host-microbiota interaction and molecular targets for therapeutic interventions.

## Acknowledgements

We thank C. Khosla and Stanford ChEM-H for use of the LC/MS qTOF instrument; A, Shiver, K.C. Huang, and A. Cheng for sharing bacterial strains; T. Meyers and T. Cowan for sharing chemical standards; J.K. Yang for consultation on web design tools; and T. Le for discussion on mass spectrometry methods. This work is supported by R01-DK085025, DP1-AT009892, R01-DK101674, generous gifts from Meredith and John Pasquesi, Heather Buhr and Jon Feiber, and a Stanford Discovery Innovation Fund Award (J.L.S.), and Chan Zuckerberg Biohub (M.A.F. and J.L.S.), DP1-DK113598 and P01-HL147823 (M.A.F.), K08DK110335 (D.D.), Stanford Dean’s Postdoctoral Fellowship and NRSA F32AG062119 (S.H.), NSF-GRFP DGE-114747 (W.V.T.), and 5T32AI007328-32 (L.G.).

## Author Contributions

S.H., W.V.T., D.D., M.A.F., and J.L.S. designed this study. S.H., W.V.T., and S.K.H. performed all bacterial culture and gnotobiotic mouse experiments. S.H. constructed mass spectrometry qTOF and QE mz-RT and ms2 spectra libraries, performed mass spectrometry experimental validation and MSDIAL data analysis, conducted comparative genomics and metabolomic distance analyses, constructed *in vivo* metabolomics pipeline and database, and designed the Metabolomics Data Explorer. W.V.T. constructed the bacterial strain library, performed phylogenetic analysis, built the *in vitro* bioinformatics pipeline and database, performed phylogenetic vs. metabolomic distance comparisons, and built random forest models. C.R.F. and L.G. conducted chemoinformatics analyses of reference library compounds. B.D.M. built the strain-resolved comparative genomics database. S.H., B.C.D., and J.M.S. developed qTOF and QE ms2 methods and collected data. D.D., L.A.F., and C.R.F. set up the MS1 mass spectrometry methods. S.H., D.D., and L.A.F. built the authentic compound collection and developed metabolomic sample preparation methods. All authors provided intellectual contributions. S.H., W.V.T., and J.L.S. wrote the paper, and all authors provided feedback.

## Materials and Methods

### Metabolomic pipeline construction logic

Accurate identification and analysis of diverse small molecules in complex biological samples (e.g., those present in the mammalian gut) is challenging due to a wide variety of technical factors including chemical structural diversity, matrix effects, and linearity of ion detection. To ensure our LC/MS pipeline is relevant for biological samples and that it is useful to the broader scientific community, we highlight six key points of our approach: 1) detectability of diverse chemical classes of compounds that characterize bacterial and host metabolism using three complementary analytical methods^42, 43^ (Extended Data Fig. 1d, 3a-c); 2) retention time (RT) shifts that occur in divergent matrices (e.g., culture supernatant vs. host serum) to determine whether metabolites in a biological sample could be faithfully identified using RT data from our mz-RT reference library (Extended Data Fig. 3d, e, Supplementary Table 2); 3) linearity of signal over a large range of concentrations, a prerequisite for performing sample comparisons and determining fold-change differences (Extended Data Fig. 3f, Supplementary Table 2); 4) use of MS/MS fragmentation to validate the high-abundance metabolites identified in biological samples (Extended Data Fig. 1e, Supplementary Table 2); 5) construction of an MS/MS spectra reference library of 750+ authentic standards on two distinct types of MS instrument (qTOF and Q Exactive) at multiple standard collision energies (Supplementary Table 3), enabling Level 1 confidence annotation when used in conjunction with our mz-RT reference library, and 6) implementation of our mz-RT reference library on different types of MS instruments following minimal non-linear RT correction^44^ (Extended Data Fig. 3g, Supplementary Table 4). For data analysis, we constructed an integrated pipeline combining 1) mass spectrometry analysis tools^45^ that leverage our reference library for compound identification (Extended Data Fig. 1f), with 2) a custom bioinformatics pipeline that enables computation and statistics on large datasets (Extended Data Fig. 2).

### Authentic chemical standard collection

The authentic metabolite standard collection is composed of individually curated and commercially available standards (Mass Spectrometry Metabolite Library of Standards, IROA Technologies). Individually curated metabolites (303 metabolites) were weighed (2 mg minimum) and transferred from original manufacturer’s stock bottles (e.g. Sigma, Fisher, Acros, etc) into 2 mL Eppendorf tubes and reconstituted with 50% LC-MS grade methanol to reach a stock concentration of 10 mM. Additional compounds (284 metabolites) were purchased as 10 mg stocks from MetaSci (MetaSci Custom Library). Dried power from company stock tubes were transferred (2 mg minimum) into 2 mL tubes and reconstituted with 50% methanol to reach 10 mM. Metabolites from the IROA metabolite standard library (634 metabolites), supplied in much smaller amounts (∼ 5 μg per well), were reconstituted with various amount of methanol in water (v/v) per manufacturer’s instructions, but due to limited mass, their concentrations were less precise. Individual pools (12-30) of metabolite standards, which do not share the same molecular weight, were generated by combining stocks and diluted with 50% MeOH to reach a final concentration of 200 μM. A subset of these pools (377 metabolites) was also serially diluted in 50% methanol. Individual metabolite pools and dilutions were analyzed using three LC/MS analytical methods.

### Mass spectrometry LC/MS methods

#### Instrumental and chromatographic settings

Compounds were separated using an Agilent 1290 Infinity II UPLC (binary pumps) and detected using an Agilent 6545 LC/MS Quadrupole Time-of-Flight (qTOF) instrument equipped with a dual jet stream electrospray ionization source (ESI) operating under extended dynamic range (EDR 1700 m/z) in the positive (ESI+) or negative (ESI-) ionization modes. Published C18 methods^42^ and HILIC method^43^ were used with minor modifications. See Supplementary Methods for details.

#### Metabolomics sample preparation

Five different sample types were processed with a similar sample preparation protocol detailed in Supplementary Methods. In brief, samples were homogenized, protein was precipitated in a methanol-based recovery buffer that contains extraction standards. Samples were centrifuged, and their supernatant was collected, evaporated, and a reconstitution buffer containing internal standards was added. Reconstituted samples were filtered and subsequently analyzed by three analytical methods on the LC/MS-qTOF.

### Mass spectrometry mz-RT reference library

The exact mass-to-charge ratio (m/z) of each metabolite standard was calculated by combining monoisotopic mass of the metabolite (PubChem) and adding or subtracting the mass of a proton (1.007276 Da) depending on the default adduct ion ([M+H]+ for ESI+ and [M-H]- for ESI-). The Agilent MassHunter Qualitative Data Analysis software (Qual, v. B.07.00) was used to match individual extracted-ion chromatogram (EIC) peaks within a ± 10 ppm window from the predicted m/z of each metabolite standard. Alternative adducts ions were identified via “Search by Molecular Feature” in Qual; when multiple adducts were identified, the adduct ion with the greatest area under the curve was used in the reference library. An RT was assigned to a metabolite when a single EIC peak was identified. When multiple chromatographic peaks were identified, likely resulting from degradation products, different isotopes, or adducts of other molecules in the mixture, a subsequent injection of that metabolite standard alone was conducted to identify the RT for that metabolite. For metabolites run in dilution series, RTs at all concentrations at which the same metabolite was detected were used to produce an averaged RT for this metabolite in the reference library. The averaged RT was used to 1) increase accuracy by averaging small injection to injection variations, and 2) distinguish true signal from background noise by validating peaks whose ion counts proportionally increase with concentrations.

To address how the same reference library performed on different instruments, we compared two different LC-MS systems: an Agilent 6545 LC-qTOF, the instrument where the original library was constructed, and a second instrument, an Agilent 6530 LC-qTOF or a Thermo orbitrap Q Exactive (QE). While these different instruments shared the same chromatographic conditions (i.e., analytical methods, solvents, columns), they differed in resolution and ESI ion source parameters optimized to support each instrument. To compare inter-instrumental RT shifts, a subset of the full reference library (219 metabolite standards spanning diverse RTs) was reconstructed on the second qTOF instrument, and 773 metabolite standards were reconstructed on the QE instrument. For each analytical method, RT correction was done by cubic polynomial transformation of the original library^44^ based on inter-instrumental RT shifts of 10-20 robustly detected metabolites (i.e., internal standards) that span the detected RT range. For each analytical method, using the corrected library with a RT tolerance window of 0.2 min, ∼99% for the 219 metabolites tested on the second qTOF instrument, and ∼94% of the 773 metabolites tested on the QE instrument, were correctly identified.

### MS/MS spectra library construction

MS/MS raw data were collected from individual pools (12-24 compounds/pool) for 833 authentic library standards, using three liquid chromatography methods applied to two distinct types of mass spectrometry instruments (Agilent qTOF 6545, Thermo Orbitrap Q Exactive (QE)). For qTOF, auto MS/MS preferred ions settings with individual input list of m/z and RT information specific to the compounds in each pool were used to collect spectra at three collision energies (CEs: 10, 20, 40 eVs). For QE, Full MS / dd-MS^2^ settings with a single shared inclusion list containing m/z and RT information for all the compound pools were used for data collection at the stepped normalized collision energy of 20-30-40%. A scan range of 60 to 900 m/z was used to collect centroid type data. On both instruments, ± 0.5 minute was used as an RT search window for MS1 peak selection, based on RT provided by the qTOF reference library. Accurate mass windows were ± 10 ppm on both instruments. RTs identified during the MS1 peak selection for the 773 compounds detected on the QE instrument were reported in the mz-RT library for the “QE_rt” column (Supplementary Table 1).

MS/MS spectra were extracted from MS/MS raw data files (mzml format) using an automated Python script (extract_ms2_spectra.ipynb) via the pymzML parsing library^46^. For each compound, the intensity of each spectral fragment was normalized to the fragment with the highest intensity (set to 1000). Spectral fragments with intensities below 0.5% relative to the highest intensity fragment were filtered out. Compound metadata (e.g., InChIKey, collision energy) and fragmentation information (e.g., m/z, intensity) were reported for each compound. Spectra from the same compound collected using different analytical methods (e.g., C18 positive and C18 negative) are all reported. In limited instances, spectra from the same compound were collected multiple times due to representation in multiple compound pools. All of the information above was compiled in Supplementary Table 3, and are publicly available at the MoNA spectra database under query phrase “Sonnenburg Lab MS2 library”. In summary, spectra from 750 and 773 unique compounds were collected on the qTOF and QE instrument, respectively.

### Mass spectrometry experimental validations

#### Linear dynamic range

For large-scale metabolomics experiments, it is typically assumed that instrument response varies linearly with analyte concentration. To test concentration linearity objectively, we constructed dilution series of 377 metabolites (from pools generated as above), in 3-fold serial dilutions spanning five orders of magnitude (from 1 nM to 200 μM). These diluted compound pools were then analyzed using the three analytical methods. Linear regression of log-transformed concentrations vs. log-transformed ion counts was performed and the coefficient of determination (*r^2^*) was calculated. Across all metabolites, the average *r^2^* and slope (on log-log plots) were both very close to 1 (0.99 and 0.92 respectively), providing a strong indication of linearity.

#### Matrix effects

The biochemical complexity of biological samples such as feces and serum may alter RT and/or detected signal of individual metabolites. To determine whether accurate identification was significantly affected by RT shifts in multiple matrices, we spiked in 132 metabolite standards into five distinct biological matrices (germ-free murine feces, serum, urine, human charcoal-stripped serum, and mega medium) and a library control condition (50% MeOH, v/v) at a final concentration of 10 μM, analyzed each matrix using all three analytical methods. Three biological replicates for each matrix were used, and RT and ion count for each spiked-in metabolite standard in each of these matrices were determined. The difference in RT between a biological matrix and the library control condition was calculated (50% methanol in water v/v) for individual spike-in metabolites. For all 132 of the metabolites in all five matrices, differences in RT were minimal, falling within a conservative ± 0.1 minute window. Changes in total ion count (area under the curve) between a biological matrix and the library control condition were determined by first removing matrix-specific, background ion counts for a small number of metabolites present in specific matrix prior to spike-in. Next, the ratio between spike-in metabolite ion counts in biological matrices and those in library blank controls was calculated (relative fold change, log2 transformed). The majority of spiked-in metabolites exhibit less than 4-fold change in ion counts relative to those detected at the library control condition (97% in murine feces, 83% in serum, 95% in urine, 88% in human serum, and 71% in mega medium). See code details in “calculate_biological_matrix_effect.ipynb”. The relatively minor impact of different biological matrices on RTs of the reference library metabolites helped establish the identification parameter (± 0.1 min RT window) for our subsequent biological experiments.

#### MS/MS validation

To verify the accuracy of compound identification obtained by our MS1 mz-RT library built from authentic standards, we unbiasedly searched MS/MS spectra of mz-RT matched individual metabolites against the Mass Bank of North America (MoNA) spectra database. MoNA-reported similarity scores based on spectra comparisons were recorded (Supplementary Table 2). For each analytical method, using the qTOF’s auto MS/MS preferred ions settings, MS/MS spectra were generated at three collision energies (CEs:10, 20, 40 eVs) from MS1 peaks identified by m/z and RT from our reference library. For biological samples, MS/MS spectra were collected for 162 high-abundance metabolites identified in QC samples from *in vitro* (bacterial supernatants) and *in vivo* experiments (*Bt*- and *Cs*-mono-associated murine samples: serum, urine, fecal/cecal contents). QC samples were generated on a per-experiment basis, by pooling equal volume from each biological replicate from the same experiment (3-8 biological replicates per condition across the entire 96-well plate) to provide a representation of the highest number of metabolites in that experiment. To establish a baseline of MoNA similarity scores, MS/MS spectra were also collected from a corresponding set of library authentic standards.

MS/MS spectra were extracted using an automated Python script by first extracting MS/MS spectra for individual mz-RT matched metabolites using pyMZML^46^, and next searching individual extracted spectra against the MoNA spectra database. The search results were restricted to spectra generated using 1) LC/MS instruments, and 2) ESI+ ionization mode (for C18 positive and HILIC positive spectra) or ESI- ionization mode (for C18 negative spectra). Each spectral search used the MoNA-default similarity score threshold of 500, and returned the top five matches with the highest similarity scores computed by the built-in MoNA algorithm. Among these top matches, the highest similarity score with the correct metabolite name was recorded (Supplementary Table 2). Because MoNA search results contained data from various LC/MS instrument platforms such as qTOF, Orbitrap, and Triple-Quadrupole, in some cases there are data collected from multiple MS platforms or multiple CEs, we would opt for the qTOF and a similar CE to our search spectra. Each MS/MS spectral comparison corresponding to the recorded score was also manually inspected. For individual metabolites repeatedly detected in the same sample type (e.g., bacterial supernatant, feces) in more than one experiment, an averaged similarity score among MS/MS spectra for the same metabolite was calculated and recorded in the summary table (Supplementary Table 2). Collectively, all similarity scores between our MS/MS spectra and MoNA spectra for the same set of metabolites have a median score of 992 (library standards, s.d. = 36.78) and 923 (biological samples, s.d. = 114) relative to a perfect score of 1000, indicating good agreement between our data and what has been previously reported.

### Data analysis

#### MS-DIAL analysis

The MS-DIAL software^45^ (v. 3.83) was used for analyzing all *in vitro* and *in vivo* data on a per-experimental run and per-analytical method basis. QC samples from each experimental run were used for peak alignment. Chemical assignment of molecular features in samples was performed by comparison of recorded RT and m/z information to our reference library constructed from authentic standards. Tolerance windows were set to 0.1 minute RT and 0.01 Da m/z for the C18 methods and 0.2 minute RT and 0.01 Da m/z for the HILIC method. When a large RT shift was observed in the internal standards (e.g., following an instrument repair), a library RT correction was done prior to MS-DIAL analysis, via polynomial transformation of the library based on inter-instrumental RT shifts of 10-20 robustly detected metabolites (e.g., internal standards). RT The minimal peak count (height) filter was set to 3000 for all experiments except for select experiments in which the MS exhibited reduced sensitivity. The MS-DIAL analysis generated a list of m/z, RT, and ion counts (area under the curve) for high-confidence annotations (matched to the reference library) as well as unknown molecular features. Based on the list of annotations for each experiment, each set of aligned peaks was manually checked using the MS-DIAL graphical user interface. Select metabolite features were removed from this list when: 1) two adjacent but distinct peaks were concurrently assigned to a single molecular feature, 2) odd curvature/shape of the peak led to integration of several “peaks” from separate sections of the same peak, or 3) features were only detected in blank controls. Annotated peaks that passed this inspection were reported in the final output file.

#### Custom bioinformatics

After MS-DIAL analysis, data were analyzed with a set of custom bioinformatics pipelines. In short, these pipelines implemented a set of filtration and normalization procedures with the goal of reducing technical variability and controlling for batch effects. The pipelines, including all code for the *in vitro* and *in vivo* sample data cleaning and standardization, are detailed at length in Supplementary Methods.

#### Distance calculations and classifiers

Comparisons between metabolomic and phylogenetic distance (Fig. 2a, Extended Data Fig. 7), and metabolite-based classification (Fig. 3a, Extended Data Fig. 8a) were done with custom Python code detailed in Supplementary Methods. For all these analyses, the metabolomic distance matrix used Euclidean distance generated from log2 transformed, media blank, delta, and variance filtered fold change data. Only the 158 strains that grew in mega medium were used for these analyses to prevent conflation of metabolic and starting medium differences.

### Bacterial culture

The bacterial strains and associated metadata (taxonomy, original repository, 16S sequence, etc.) used in this work are recorded in Supplementary Table 6. All bacterial inoculation and growth occurred in a Coy Laboratories anaerobic chamber kept at an atmosphere of approximately 80:15:5 (N_2_: CO_2_: H_2_, percent). All incubations occurred at 37**°**C, all bacterial stocks were stored at -80**°**C, and all optical densities (ODs) were recorded at 600nm using a BioTek Epoch 2 plate reader.

#### Stock preparation

Bacterial strains were acquired from various culture collections including ATCC, DSMZ, NCTC, and BEI. Source cultures were plated on a rich medium, single colonies picked, cultured in rich medium, and stored as 1mL frozen cultures (25:25:50 v/v glycerol:H_2_O:culture) in ThermoFisher Matrix Tubes. The solid and liquid media used for stock generation are recorded in Supplementary Table 6 (worksheet ‘media’). Source cultures that exhibited multiple morphologies on agar plates were purified and morphologies separated and retained if the 16S sequence matched the expected 16S sequence. For all cultures, the purity of the final cultures was checked by 16S rRNA sequencing (see Supplementary Methods).

#### Bacterial media

All media used in this study are recorded in Supplementary Table 6 (worksheet ‘media’). Note that in some cases we grew and recorded metabolites from taxa in multiple media. For the media used for particular supernatant samples and metabolomics see Supplementary Table 7 (worksheet ‘aggregated_md’).

Mega medium was prepared according to the protocol in Supplementary Methods. The recipe is slightly adapted from a previous publication^47^. In our usage of mega medium, each batch was autoclaved, moved into the anaerobic chamber, and allowed to become anaerobic for at least 24 hours before use. For taxa that would not grow in mega medium, a different medium was selected based on literature. In each case, we referenced an ATCC, DSMZ, or media manufacturer (e.g., Hardy Diagnostics) recipe as outlined in Supplementary Table 6 (worksheet ‘media’). In all cases, these media were prepared for use similarly to mega medium. Specifically, adjustment of pH was done prior to autoclave, filter sterilized vitamins and sterile blood were added after autoclave, and media was moved immediately from autoclave to anaerobic chamber and allowed to become fully anaerobic for at least 24 hours prior to use.

For identification of nitrogen utilization in Bacteroidetes, Salyer’s minimal medium (SMM) was prepared (see Supplementary Methods) slightly modified from published protocols^26, 48^. In short, SMM base was prepared (SMM without hematin, nitrogen source, or reduced sulfur source) and allowed to become anaerobic in foil-covered bottles. SMM was prepared without nitrogen source to avoid spontaneous glutamine degradation^49^. Immediately prior to use, SMM base was amended with filter sterilized solutions of hematin (final concentration 0.5mg/100ml), nitrogen source (glutamine, asparagine, glutamic acid, or ammonium sulfate, final concentration 10 mM), and reduced sulfur source (cysteine or sodium sulfide, final concentration 4.12 mM). Taxa were plated (mega medium or brain heart infusion with blood) and a single colony picked into freshly prepared SMM. Preculture for 24 hours was followed by subculture into freshly prepared SMM for 12-36 hours. OD readings were taken as above.

#### In vitro growth for metabolomics

Bacterial supernatants included in the *in vitro* data were generated according to the following protocol. Cultures were inoculated into anaerobic medium (∼4 μL: 1600 μL) in triplicate in 2 mL 96-well blocks and incubated for 24-72 hours depending on the taxa selected. Therefore, a single biological replicate from the bacterial culture experiments represents an individual well or tube of bacterial culture growth from an independent 4 uL aliquot from a frozen glycerol culture stock. These pre-cultures were subcultured into mega medium (∼4 μL: 1600 μL) and similarly incubated for 12-60 hours. 200 μL of subculture was incubated in a plate reader so that OD readings could be taken to monitor growth phase. The remaining cell cultures were harvested when OD readings showed late log or early stationary phase. Harvested culture was immediately removed from the anaerobic chamber, centrifuged to pellet cells (5,000 g, 10 min), and cell-free supernatant was either frozen at -80**°**C or immediately extracted as described in Supplementary Methods.

For purity analysis, sequencing protocol, and phylogenetic tree reconstruction see Supplementary Methods for details.

### Mouse experiments

Mouse experiments were performed on gnotobiotic Swiss Webster germ-free mice (males, 10-14 weeks of age, n = 3-8 per group for all experiments) or Swiss-Webster Excluded Flora mice (“conventional mice”, males, 10-14 weeks of age, n = 3 per group) maintained in aseptic isolators, and originally obtained from Taconic Bioscience. Mice were maintained on a 12-hour light/dark cycle at 69 °F in ambient humidity, fed ad libitum, and maintained in flexible film gnotobiotic isolators for the duration of all experiments (Class Biologically Clean, Madison WI). For mono-association experiments, mice were colonized with *Bacteroides thetaiotaomicron* VPI 5482*, Clostridium sporogenes* ATCC 15579, *Citrobacter portucalensis* BEI HM-34, or *Anaerostipes sp.* BEI HM-220by oral gavage (200 uL, ∼1 x 10^7^ CFU) and were maintained on a standard chow (LabDiet 5K67). For the defined-community experiment, mice with a six-member community were colonized with a 200 uL mixture consisting of equal volumes from saturated cultures of *Bacteroides thetaiotaomicron* VPI 5482 (8.7 x 10^9^ CFU), *Clostridium sporogenes* ATCC 15579 (1.4 x 10^8^ CFU)*, Edwardsiella tarda* ATCC 23685 (3.6 x 10^10^ CFU), *Collinsella aerofaciens* ATCC 25986 (1.4 x 10^9^), *Eubacterium rectale* ATCC 33656 (6.9 x 10^6^ CFU), and *Parabacteroides distasonis* ATCC 8503 (1.5 x 10^9^ CFU). Mice with a five-member community were colonized with all cultures mixed at the same volumes as described above except for *Clostridium sporogenes* ATCC 15579, which was not included. Successful colonization and stable community members were determined by 16S amplicon sequencing of the V4 (515f, 806r) region of microbial populations present in the feces and cecal contents from individual mice.

For all experiments, mice were euthanized by CO_2_ asphyxiation nine days (mono-colonization with *Citrobacter portucalensis* BEI HM-34 or *Anaerostipes sp.* BEI HM-220) or four weeks (all other experiments) following colonization, and four sample types (serum, urine, feces, and cecal contents) were harvested from each mouse. A single biological replicate in the mouse experiments represents a specific sample type (e.g., serum) harvested from an individual mouse (i.e., each biological replicate is from a different mouse). Prior to euthanization, urine and feces were collected. Whole blood was collected by cardiac puncture and serum was obtained using microcontainer serum separator tubes from Becton Dickinson following manufacturer’s instructions. The intact cecum was harvested and snap-frozen in liquid nitrogen. A single cecal sample was harvested for mono-association and conventional experiments, and three samples at three different sections of the cecum were harvested for the defined-community experiment. All mouse experiments were conducted under a protocol approved by the Stanford University Institutional Animal Care and Use Committee.

### Comparative genomics

#### Genome annotation and database

Bacterial isolates from the culture collection were manually linked up to their respective NCBI BioProject ID numbers. The Rentrez package (https://cran.r-project.org/package=rentrez) was used to link BioProject ID numbers with existing GenBank or RefSeq assemblies or with reads from the Sequence Read Archive (SRA) for isolates which were previously sequenced but not assembled. Isolates lacking assembly accession numbers Supplementary Table 6 (worksheet ‘full_taxonomy’) were assembled using previously described methods^50^. Briefly, reads were trimmed using Trimmomatic^51^ and assembled using SPAdes v3.9.1^52^ using the following parameters: k=21,33,55 --careful --cov-cutoff auto. Contigs smaller than 1500bp were removed, and assemblies were gene-called and annotated using prokka v1.14.5^53^. MultiGeneBlast^54^ (v1.1.13) was used to build a database containing all assembled and downloaded genomes listed in Supplementary Table 6.

#### Gene and gene cluster searches

The *arc* gene cluster from *Lactococcus lactis* and the *spe* gene cluster from *Escherichia coli* were used as the query to search publicly available, assembled genomes of strains within our collection. Comparative genomics analyses were conducted using the “Architecture Search” feature of the MultiGeneBlast software (v. 1.1.13) with default parameters with one modification which set the “maximum distance between genes in locus (kb)” to 40 kb. For identification of Asparaginase-containing genomes, the custom BLAST database described above was queried for homologs to *E. coil genes* (*ansA, ansB*, and *aspA*) that encode asparagine-consuming enzymes.

### Metabolomics Data Explorer

The Metabolomics Data Explorer (https://sonnenburglab.github.io/Metabolomics_Data_Explorer) was constructed in JavaScript and generates scatter plots of our *in vitro* and *in vivo* fold-change data based on user input. *In vitro* and *in vivo* metadata and fold-change data files are used as data input and are parsed using the Papa Parse library to extract the data and populate the dropdown menus on each page. The dropdown menus enable users to pick the desired taxonomy, metabolite, media (*in vitro*), and colonization, metabolite, sample type (*in vivo*). The Nivo library is used to render interactive scatter plots of the fold change data relative to media blank controls (*in vitro*) or to germ-free controls (*in vivo*). Each dot represents an independent biological replicate, and all metabolites (uniquely identified or co-eluting) are shown. In rare cases, the same metabolite may appear twice in the scatter plot if it is uniquely identified in one analytical method while co-eluting with other metabolites in another analytical method. The scatter plot presents all biological replicates from all independent experiments available in the dataset, and provides label details when hovering over data points to enable easy identification.

### Data availability

All metabolomics raw data are publicly available on the Metabolomics Workbench under study number ST001683 for *in vivo* data and study number ST001688 for *in vitro* data. MS/MS spectra libraries generated on the qTOF and QE instruments are publicly accessible on the MoNA spectra database (https://mona.fiehnlab.ucdavis.edu), and can be queried using keywords “Sonnenburg Lab MS2 Library.”

### Code Availability

Custom Python code was written to enable the construction of the MS/MS spectra libraries, the processing and visualization of the *in vitro* and *in vivo* LC/MS data, the optical density and growth curve data, the bioinformatic analysis of 16S and whole genomes, and the analysis of the metabolomics data. Full code for each of these steps is provided at the Sonnenburg lab GitHub site (https://github.com/SonnenburgLab/Han_and_Van_Treuren_et_al_2021). The JavaScript code supporting the interactive, web-based Metabolomics Data Explorer is also provided (https://github.com/SonnenburgLab/Metabolomics_Data_Explorer)

## Extended Data Figure Legends

**Extended Data Fig. 1.**
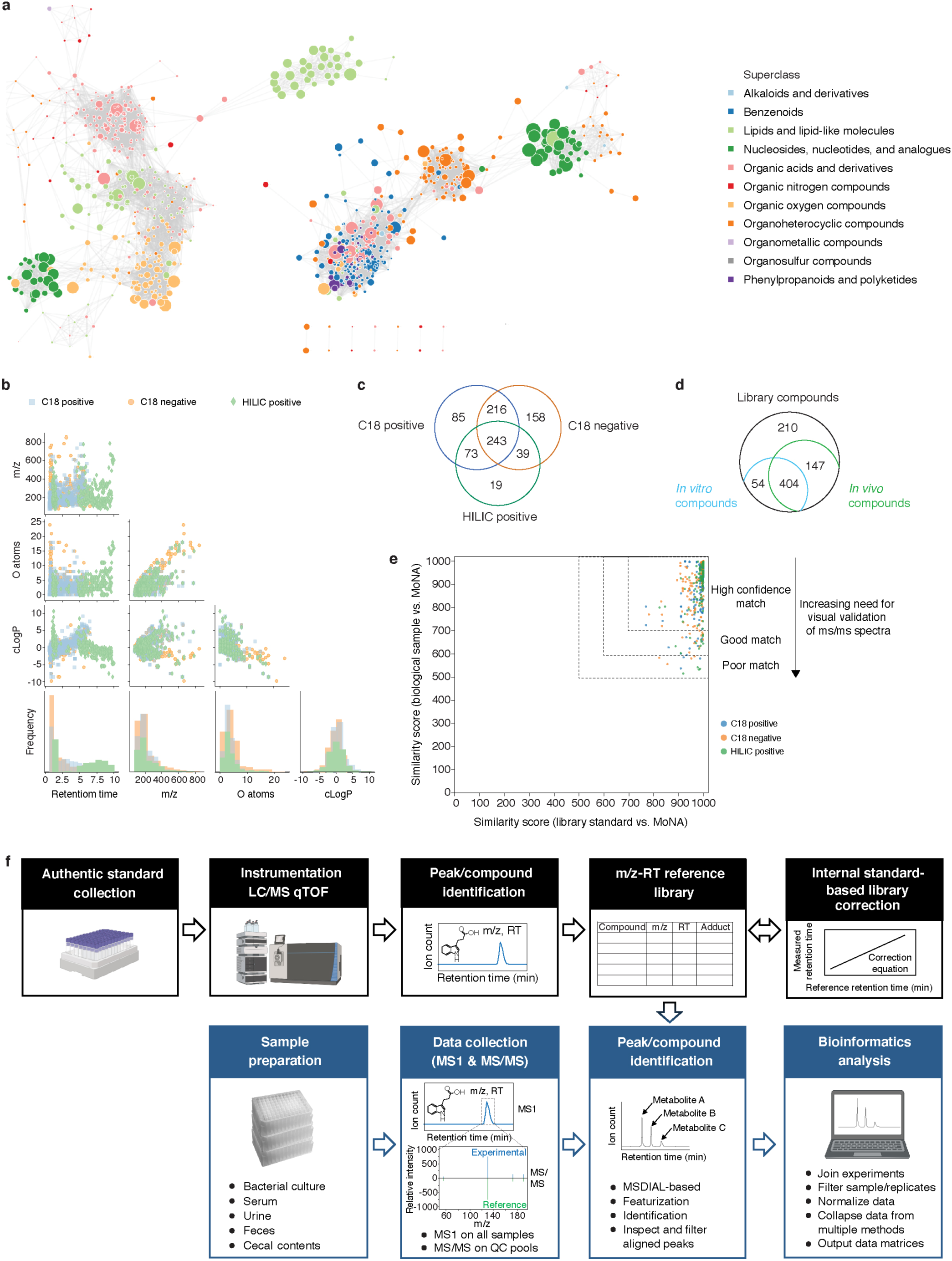
Summary statistics on mass spectrometry reference library metabolites, their detection, and validation. **a,** Chemical similarity network of the compound library. Network nodes: library compounds colored by their superclasses. Node size: monoisotopic mass. Edges between nodes: substructure similarity values above a z-score threshold of 1 standard deviation from the mean. **b,** Scatter plots and histograms of chemical properties of 833 library metabolites. **c,** Venn diagram of library compounds that are detected by each of the three methods. **d,** Venn diagram of compounds (by PubChem CID) identified in the reference compound library (Supplementary Table 1), *in vitro* conditions (Supplementary Table 7, “count.ps”), and *in vivo* conditions (Supplementary Table 8, “istd_corr_ion_count_matrix”). *In vitro* conditions include all media types, and *in vivo* conditions include all sample types: urine, serum, feces, and cecal contents, and all colonization states. **e,** Scatterplot of all pairwise similarity scores (biological sample vs. library) of the same compound searched against the MoNA spectra database. All library standards (median similarity score = 992) and 97.3% of corresponding compounds from biological samples (median similarity score = 923) exhibit similarity scores ≥ 600, and 2.7% of those compounds from biological samples score below 600. Confidence levels are determined based on both similarity scores and visual validation of the MS/MS spectra. **f,** Schematic of the metabolomics pipeline’s data collection and analysis workflow.

**Extended Data Fig. 2.**
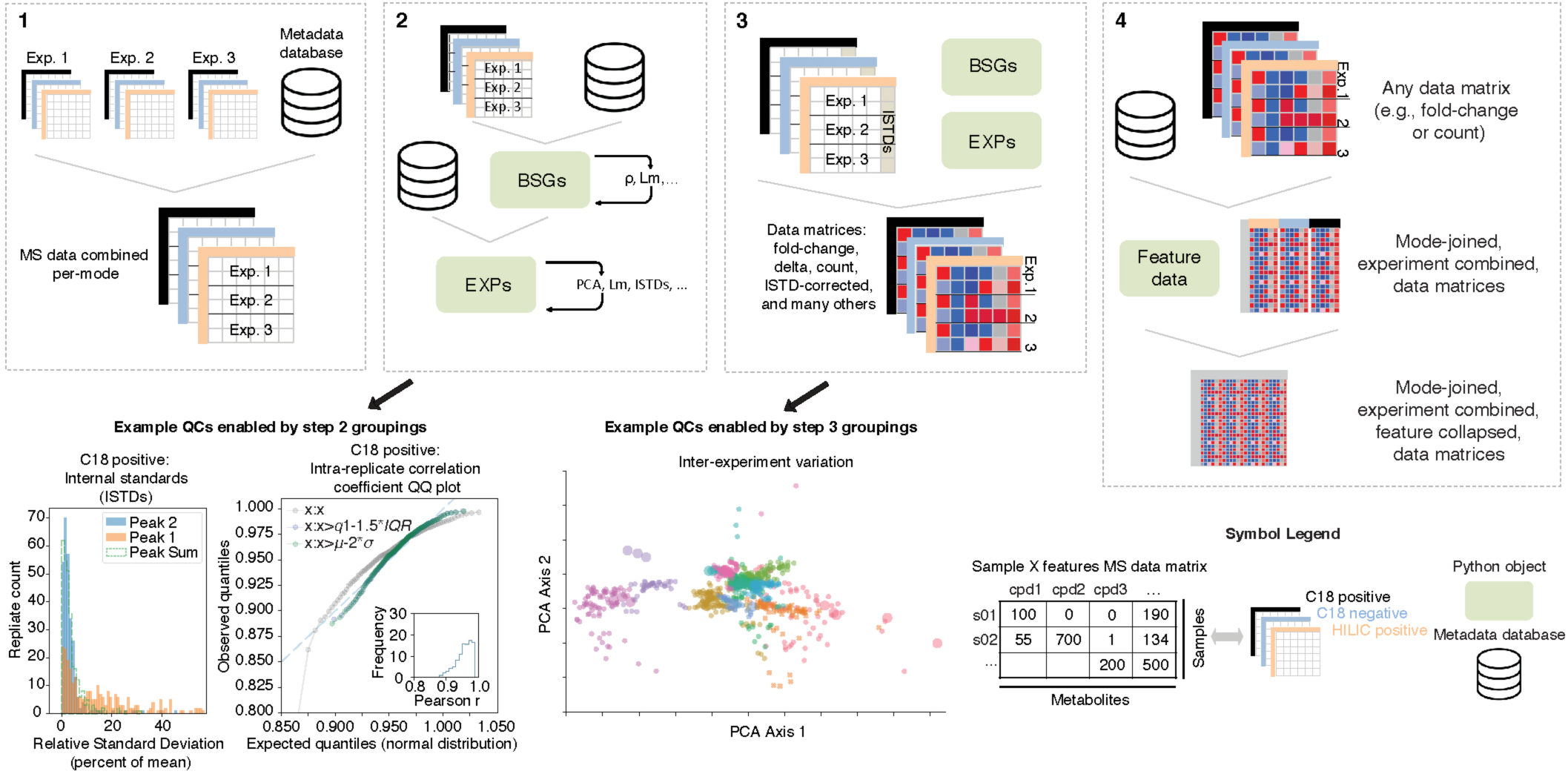
Schematic of a custom bioinformatics analysis pipeline that generates a metabolite fold-change matrix. The pipeline integrates data across multiple experimental runs and minimizes intra-replicate, intra-experiment, and inter-experiment variability. The four steps detailed here are explained in depth in the Supplementary Methods (*Custom bioinformatics: in vitro pipeline*). Step 1: A database recording sample metadata (organism, media, growth data, etc.) and MS-DIAL output files are integrated into data matrices specific to each analytical method. Step 2: All data are grouped by replicate (Biological Sample Groups; BSGs) and analyzed to remove replicates with low intra-replicate correlation. Replicates are then grouped by experiment (EXPs) to assess inter-experiment variability. Transformations reducing inter-experiment variability are identified and compared. For metabolites that are detected by multiple methods, their ion counts are compared on a per-replicate and per-experiment basis to identify one or more methods that consistently detect these metabolites. Step 3: Using an internal standard-based correction, ion counts for individual samples are adjusted and transformed into different fold-change data matrices. Step 4: Data matrices corresponding to each method are combined into a single data matrix representing all detected metabolites.

**Extended Data Fig. 3.**
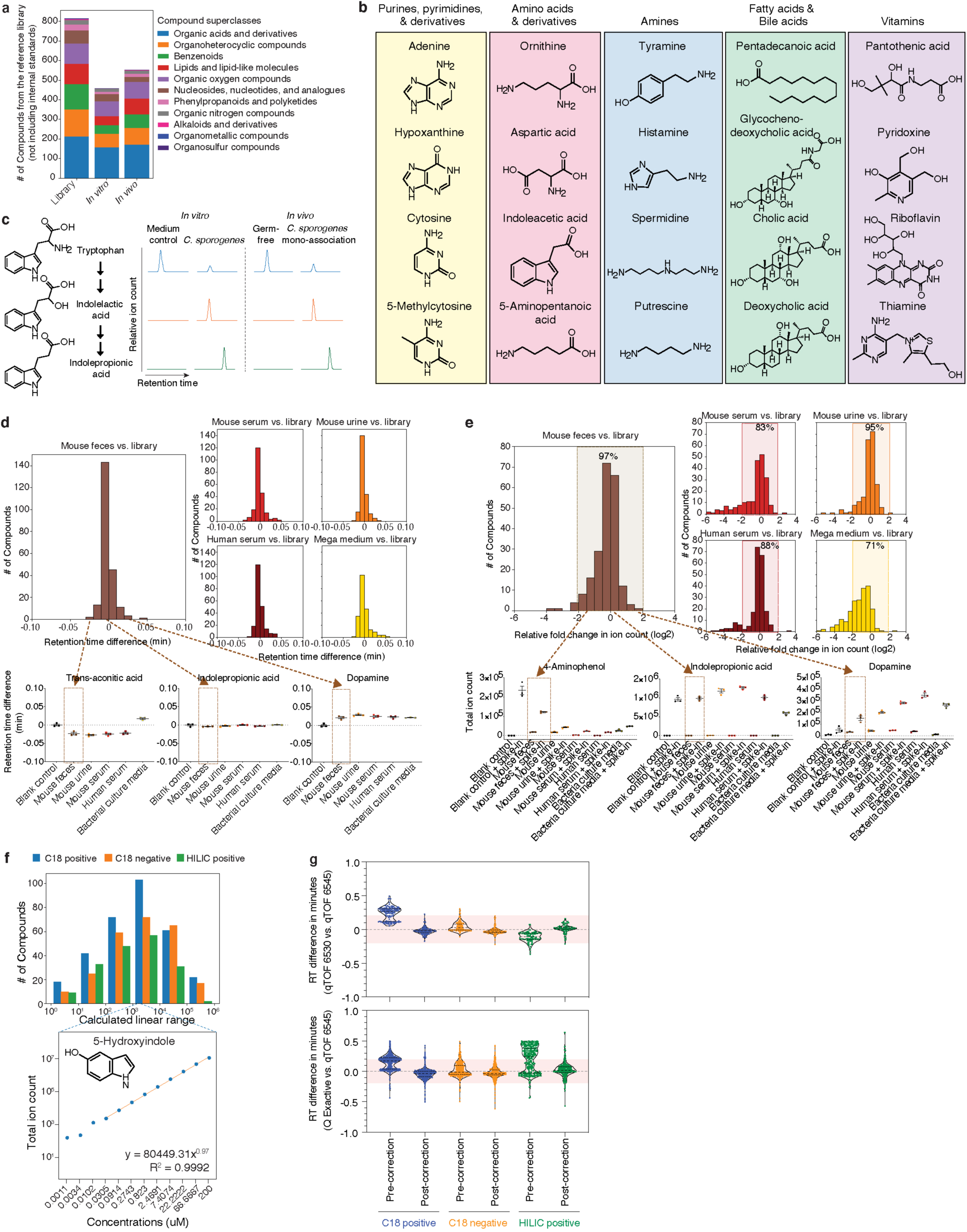
High-throughput identification and analysis of diverse metabolites in complex biological matrices. **a,** Number of unique compounds (by PubChem CID) within distinct chemical superclasses detected in the mz-RT reference library (n = 815, 11 superclasses), *in vitro* dataset (n = 458, 9 superclasses), or *in vivo* dataset (n = 551, 9 superclasses), excluding internal standards. Nine of the 11 chemical superclasses in the reference library are represented in metabolites detected *in vitro* and *in vivo*. The two remaining library superclasses (Organosulfur and Organometallic compounds) not represented in the experimental data contain one compound each. **b,** Diverse classes of metabolites identified in the conventional murine cecum. Representative metabolites shown are significantly elevated (≥ 4-fold, corrected *P* < 0.05) in the conventional mice vs. germ-free controls in one experiment with n = 3 (conventional) and n = 4 (germ-free) mice. *P* values: two-tailed *t*-test with Benjamini-Hochberg correction for multiple comparisons. **c,** Examples of precursor, intermediate, and products from the tryptophan fermentation pathway being identified by our methods both *in vitro* (*Cs* culture supernatant) and *in vivo* (*Cs* mono-association cecal contents). Extracted ion chromatogram peaks representing relative ion counts for each metabolite are shown. **d, e,** Histograms of changes in retention time (RT) (**d**) and total ion count (**e**) for 132 spike-in metabolites in five complex biological matrices using three analytical methods. All spiked-in metabolites show minimal change in RT, falling within a conservative ± 0.1 minute search window from their RTs as determined in the library control condition (**d**). The majority of spiked-in metabolites (e.g., 97% in feces) exhibit less than 4-fold change in ion counts relative to those detected at the library control condition (**e**). Representative examples of RT shifts (**d**) and changes in total ion counts (**e**) in individual metabolites in the mouse fecal matrix are shown. Mean ± s.e.m. of one experiment with n = 3 biological replicates. ean. **f,** Histograms of linear ranges of 377 reference library metabolites measured in serial dilutions. A representative linear range of 5-Hydroxyindole is shown. **g,** Violin plots (median, quartiles) of differences in RTs measured by three analytical methods between distinct mass spectrometry instruments: qTOF 6454 where the library was built vs. a second instrument qTOF 6530 for a shared panel of 219 reference library metabolites (upper panel) or vs. a second instrument orbitrap Q Exactive for a shared panel of 773 reference library metabolites (lower panel). Mean RT differences (in minutes) between two instruments by each method (C18 positive, C18 negative, and HILIC positive, respectively): qTOF vs. qTOF, upper panel (pre-correction: 0.238, 0.044, -0.110; post-correction: -0.023, -0.020, 0.015); qTOF vs. QE, lower panel (pre-correction: 0.151, 0.027, 0.196; post-correction: -0.040, -0.021, 0.026). Per method, RT correction was performed by polynomial transformation of the library based on inter-instrumental RT shifts of 10-20 robustly detected metabolites. Per method, using the corrected library with a RT tolerance window of 0.2 min, ∼99% of the 219 metabolites tested on the second qTOF and ∼94% of the 773 metabolites tested on the QE were correctly identified.

**Extended Data Fig. 4.**
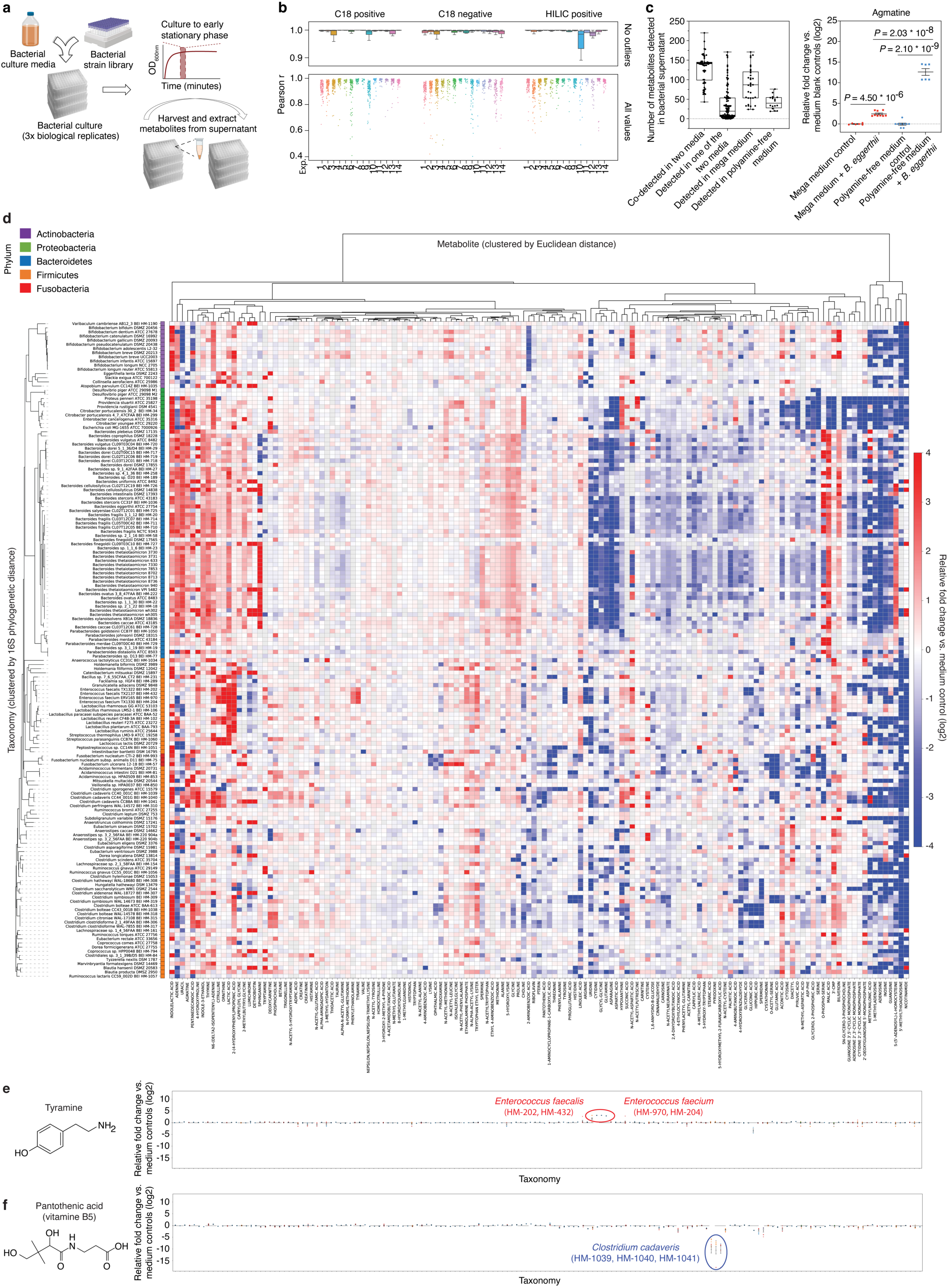
Conserved and unique metabolomic signatures across bacterial taxa. **a,** Schematic of our high-throughput bacterial culture and sample collection workflow. **b,** Intra-replicate Pearson correlation coefficients (triplicates and greater) stratified by 14 independent bacterial culture experiments and three analytical methods. For each experiment, Pearson correlation r values were calculated for all supernatant and media sample replicate groups: n = 346 (C18 positive), n = 344 (C18 negative), and n = 344 (HILIC positive). Total ion count data were corrected by internal standards and log transformed, standardized and scaled, prior to computing Pearson correlation values. Box: median, 25th, and 75th percentile; whiskers: Tukey’s method. **c,** Left panel: Number of medium-specific or common metabolites detected in the same bacterial strain grown in two different media (29 strains cultured in two or more of the 12 different media). Each dot represents the total number of metabolites from a single comparison between two media in which a strain has been grown: n = 58 (co-detected in two media), n = 116 (detected in one of the two media), n = 33 (detected in the mega medium), and n = 16 (detected in polyamine-free medium). Box: median, 25th, and 75th percentile; whiskers: minimum and maximum. Right panel: Agmatine production levels by *B. eggerthii*. Mean ± s.e.m from two to three independent experiments, each with n = 3 biological replicates. *P* values: two-tailed *t*-test with Benjamini-Hochberg correction for multiple comparisons. **d,** Heatmap of metabolomic profiles of 158 mega medium-grown bacterial strains, clustered by 16S phylogenetic distance. Individual metabolites are hierarchically-clustered (Ward’s method) using Euclidean distance between the fold-change (log2 transformed) values across all taxonomies. Metabolites shown are detected in at least 50% of the 158 taxonomies to enable Ward clustering. **e, f,** Production or consumption patterns of tyramine and pantothenic acid across 158 mega-medium grown strains. Mean ± s.e.m from one to three independent experiments (by dot color), each with n ≥ 3 biological replicates.

**Extended Data Fig. 5.**
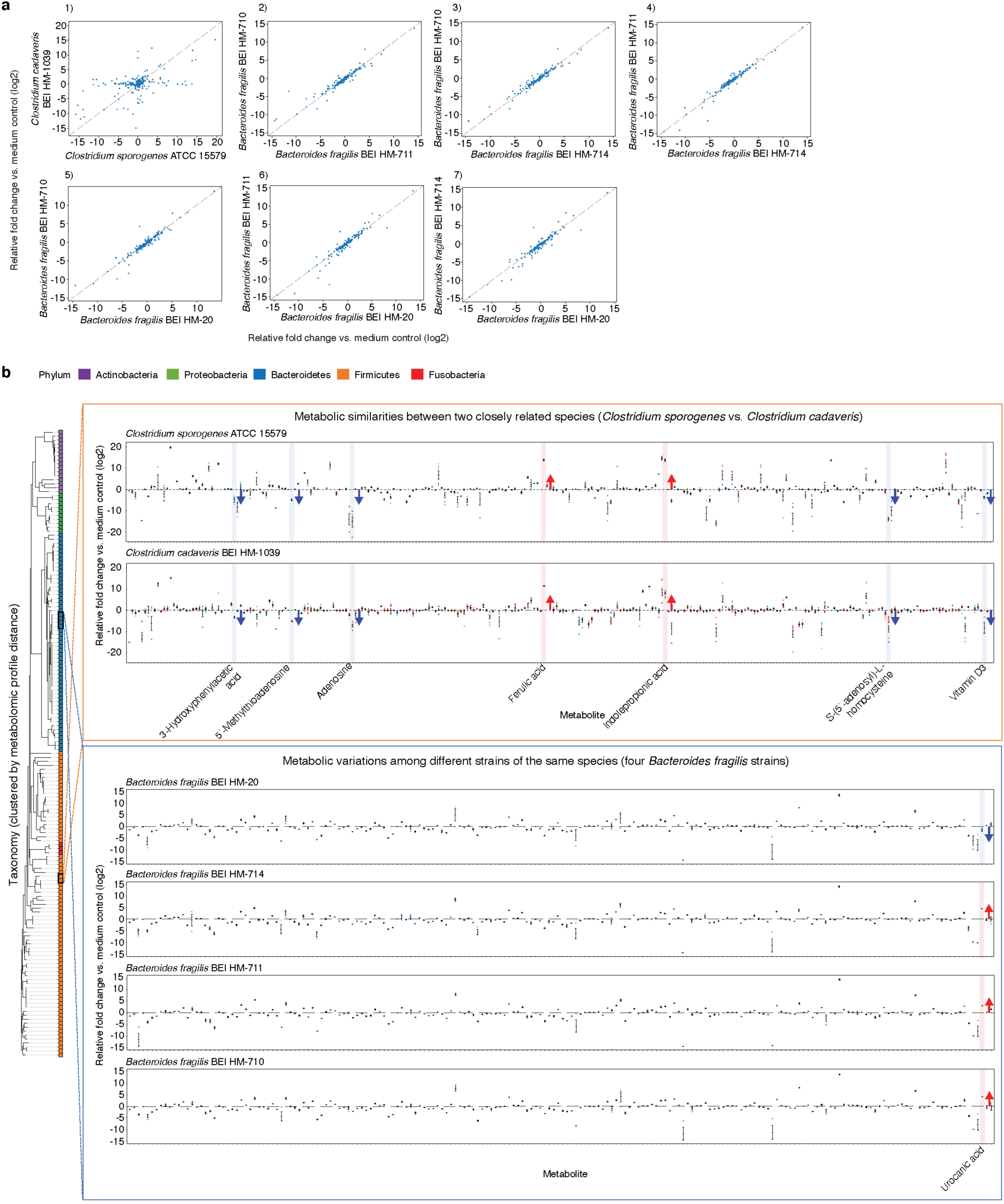
Metabolic profile variation among related bacteria. **a,** Pairwise metabolomic profile comparisons between two closely related strains grown in mega medium: *Clostridium sporogenes* ATCC 15579 and *Clostridium cadaveris* HM-1039 (subpanel 1), and among four strains of *Bacteroides fragilis* (subpanels 2-7): HM-710, HM-711, HM-714, and HM-20. Each dot represents an averaged fold-change value (log2-transformed) from one to three independent experiments, each with n = 3 biological replicates. Pearson correlation r values of pairwise metabolomic profile comparisons, performed on standardized and scaled data: ATCC 15579 vs. HM-1039 (r = 0.063), HM-711 vs HM-710 (r = 0.859), HM-714 vs. HM-710 (r = 0.866), HM-714 vs. HM-711 (r = 0.880), HM-20 vs. HM-710 (r = 0.829), HM-20 vs. HM-711 (r = 0.845), and HM-20 vs. HM-714 (r = 0.807). **b,** Metabolic similarities and variations among closely related species of *C. sporogenes* and *C. cadaveris*, and among different strains of the same species of *B. fragilis* grown in mega medium. Taxonomies shown are clustered by 16S phylogenetic distance, and are colored by distinct phyla. Mean ± s.e.m. from one to three independent experiments, each with n = 3 biological replicates

**Extended Data Fig. 6.**
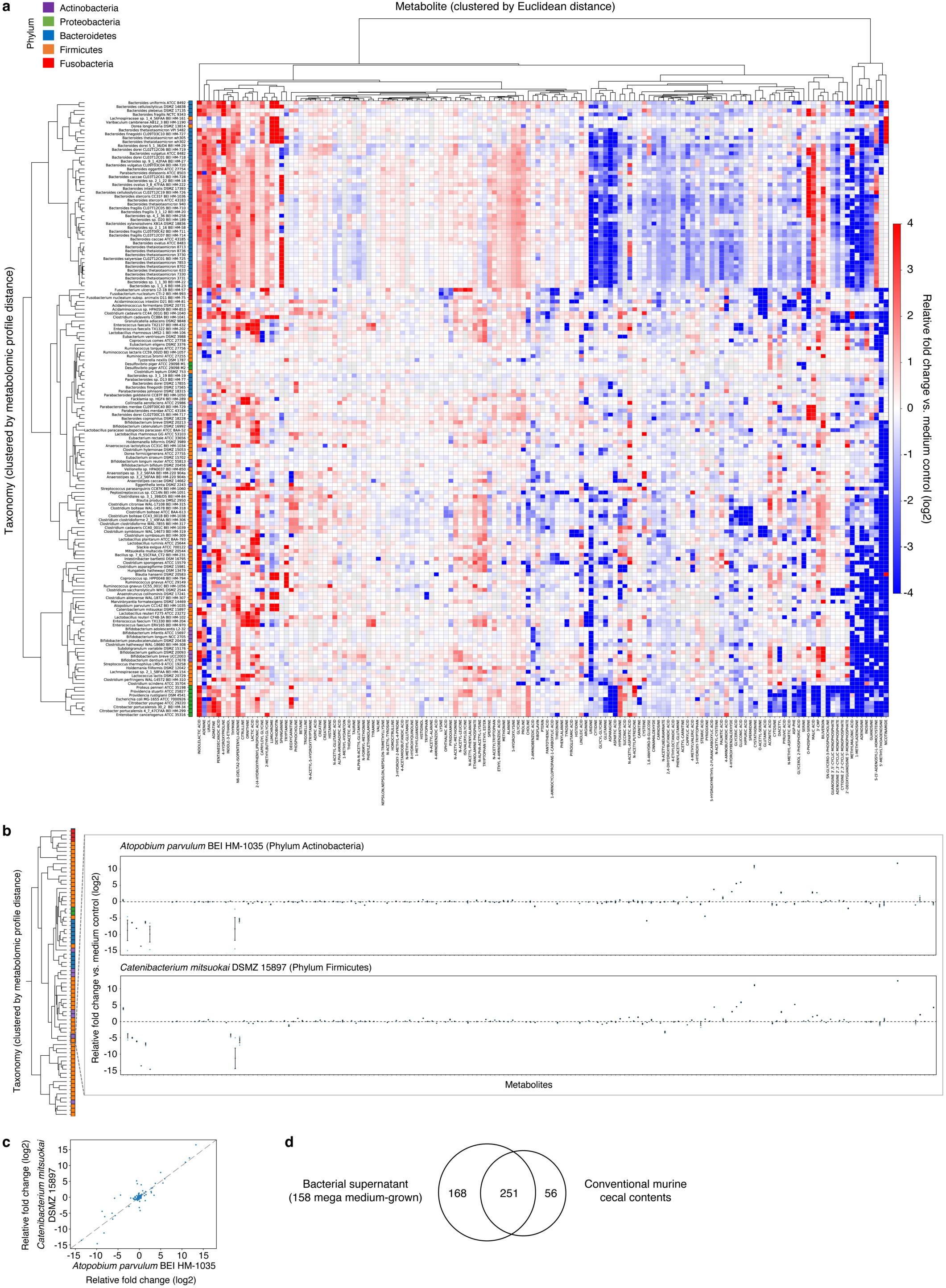
Relationships between phylogeny, taxonomy, and metabolome. **a,** Metabolomic profiles of 158 mega-medium grown bacterial strains. Individual taxonomies are clustered by metabolomic profile distances (fold change, log2 transformed) across all metabolites. Individual metabolites are hierarchically-clustered (Ward’s method) using Euclidean distance between the fold-change (log2 transformed) values across all taxonomies. Metabolites shown are detected in at least 50% of the 158 taxonomies to enable Ward clustering. **b,** Metabolic similarities between two phylogenetically distant species grown in mega medium. Taxonomies are clustered by metabolomic profile distances (fold change, log2 transformed) across all metabolites. Mean ± s.e.m. of one experiment with n = 3 biological replicates. **c**, Scatter plot of pairwise metabolomic profile comparison between two phylogenetically distant species. Each dot represents an averaged fold-change value (log2-transformed) of one experiment with n = 3 biological replicates. Pearson correlation of pairwise metabolomic profile comparison between these two species, performed on standardized and scaled fold-change data, r = 0.7090. **d**, Venn diagram of unique and overlapping compounds (by PubChem CID) identified in culture supernatant of 158 mega-medium grown strains and cecal contents of conventional mice.

**Extended Data Fig. 7.**
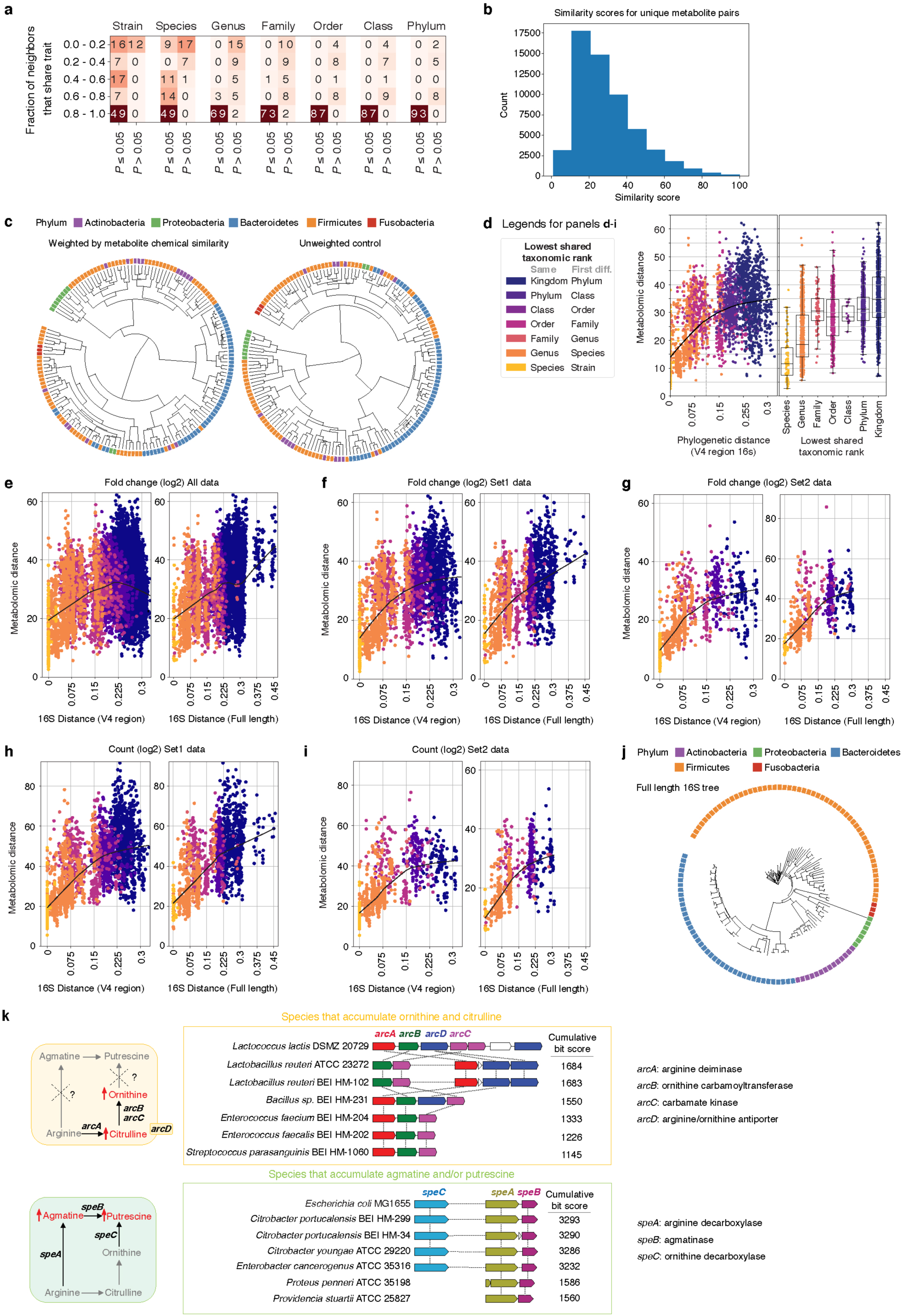
Multiple data transformations identify non-linear relationship between phylogenetic and metabolomic distance. **a,** Heatmap showing comparison of phylogenetic and metabolomic tree topologies. Cells record the number of tips whose neighborhoods share more overlap than expected (*P* < 0.05; one-sided permutation test). Data are stratified by fractional overlap of neighborhoods and permutation probability (Supplementary Methods: *Distance Comparisons*). **b,** Histogram of chemical similarity scores (based on Tanimoto 2D structures) between each unique pair of compounds (by PubChem CID) detected in the *in vitro* dataset. 359 non-coeluting compounds were used for this pairwise comparison. **c**, Metabolomic distance tree with each metabolite weighted based on their chemical similarity (left) or unweighted control metabolomic distance tree (right). The weighted and unweighted matrices were calculated using uniquely detected, non-coeluting compounds in the *in vitro* dataset, where a unique PubChem CID identifier can be assigned to each compound. Two-sided Mantel test for comparison between the weighted and unweighted distance matrices: r^2^ = 0.863, *P* = 0.001. **d,** Left panel: Correlation of phylogenetic and metabolomic distance across pairs of strains colored by lowest shared taxonomic rank with a LOESS fit shown. Right panel: Metabolomic distance between pairs of strains binned by the lowest shared taxonomic rank. Species (n = 111), Genus (n = 1386), Family (n = 159), Order (n = 1222), Class (n = 34), Phylum (n = 1442), and Kingdom (n = 8442). Box: median, 25th, and 75th percentile; whiskers: Tukey’s method. Internal standard (IS)-corrected fold-change data (**e**-**g**) and IS-corrected total ion count data (**h**, **i**) were log-transformed and used to calculate pairwise metabolomic distances between microbial taxa. These distances were compared to the corresponding pairwise phylogenetic distances generated from a tree built with the V4 region of 16S (left column) or the full-length 16S gene (right column). Data are plotted with a LOESS fit. Set1: microbes grown in at least one experiment simultaneously. Set2: microbes grown in the same experiment only. **j,** Phylogenetic tree constructed using full 16S sequences of a subset of the mega-medium grown strains. Only strains with available full 16S sequences are shown (Supplementary Table 6). **k**, Left panel: Schematic of the pathway that synthesizes citrulline and ornithine, or synthesizes agmatine and/or putrescine. Right panel: The top six matches identified by the comparative genomics tool MultiGeneBlast within a 40-kb search window, when searched against a genomic database of our strain collection with sequenced genomes. Horizontal dashed lines between genes represent multiple other genes present within the search window.

**Extended Data Fig. 8.**
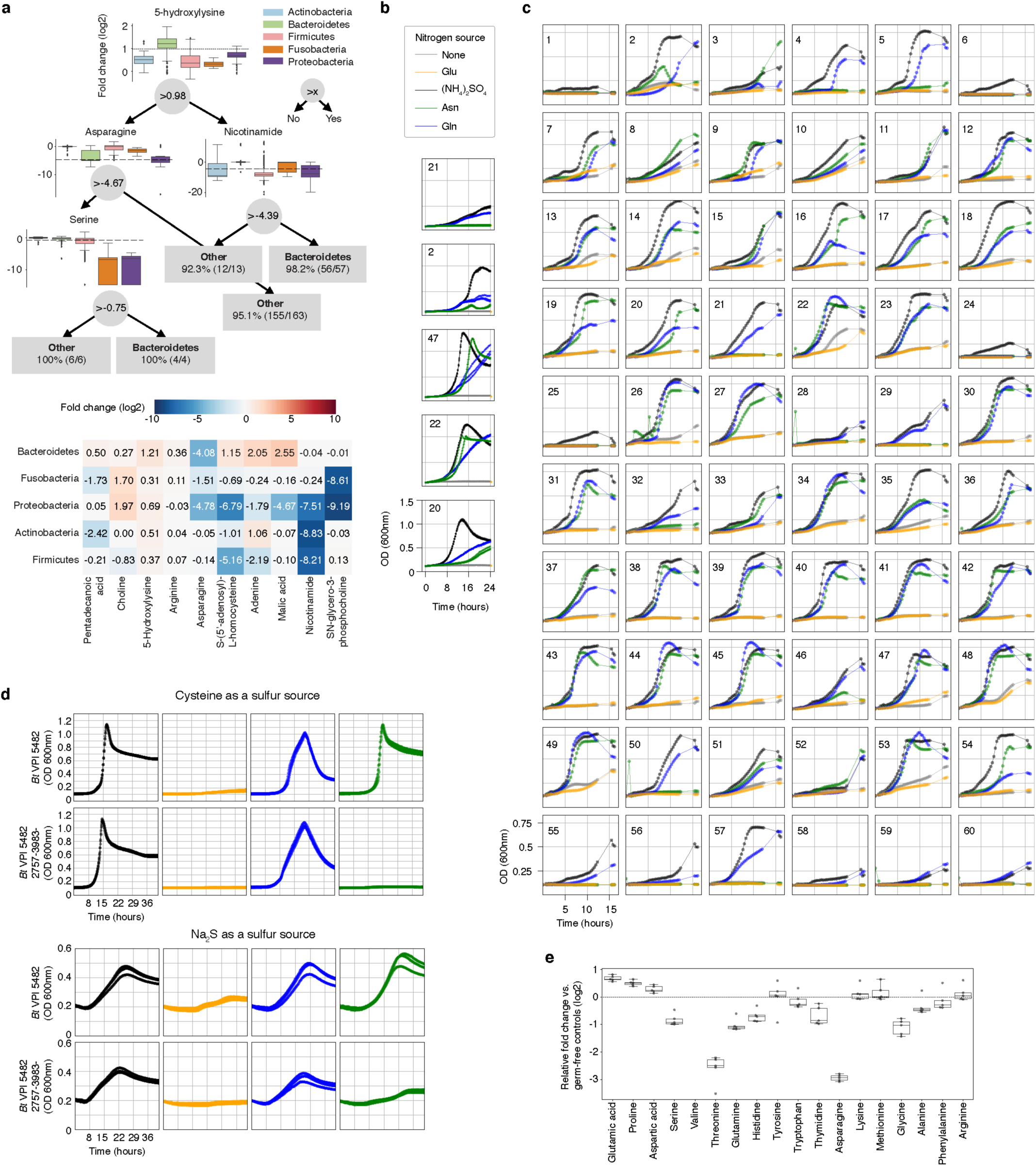
Asparagine and glutamine can be utilized as sole nitrogen sources by most tested Bacteroidetes. **a,** Top panel: An example decision tree from a forest that can differentiate Bacteroidetes vs. bacteria from the other four represented phyla with > 97% accuracy. For each decision node, phylum-level elevation and reduction based on metabolite levels are shown (relative fold change to bacterial media controls, log2 transformed). Actinobacteria (n = 20), Bacteroidetes (n = 57), Firmicutes (n = 83), Fusobacteria (n = 3), and Proteobacteria (n = 10). Dashed line: metabolite threshold. Box: median, 25th, and 75th percentile; whiskers: Tukey’s method. Bottom panel: The 10 most important features differentiating the five tested phyla. Data are shown as median metabolite log2 fold-change for each phylum; metabolites and phyla are ordered by Ward linkage distance. **b,** Representative growth curves from two independent experiments, each with n = 3 biological replicates for a subset of Bacteroides spp. using modified Salyer’s Minimal Medium (SMM) with indicated nitrogen source. Legend colors for sole nitrogen source are maintained for panels **b**-**d**. **c,** Representative growth curves of one experiment with n = 5 biological replicates for 60 Bacteroidetes using modified SMM with indicated nitrogen sources. **d,** Growth curves of wild-type and mutant *Bt* grown in defined minimal media with either cysteine (top) (one experiment, n = 3 biological replicates) or sodium sulfide (Na_2_S, bottom) as sole reduced sulfur sources (one experiment, n = 3 biological replicates). **e,** Amino acid production and consumption levels in gnotobiotic mice mono-associated with *Bacteroides thetaiotaomicron* (*Bt*) (one experiment, n = 5 mice). Box: median, 25th, and 75th percentile; whiskers: Tukey’s method). Numeric labels in **b** and **c** correspond to the following: 1: *B. acidifaciens* DSMZ 15896, 2: *B. caccae* ATCC 43185, 3: *B. caccae* BEI HM-728, 4: *B. cellulosilyticus* BEI HM-726, 5: *B. cellulosilyticus* DSMZ 14838, 6: *B. coprophilus* DSMZ 18228, 7: *B. dorei* BEI HM-29, 8: *B. dorei* BEI HM-717, 9: *B. dorei* BEI HM-718, 10: *B. dorei* BEI HM-719, 11: *B. dorei* DSMZ 17855, 12: *B. eggerthii* ATCC 27754, 13: *B. eggerthii* DSMZ 20697, 14: *B. finegoldii* BEI HM-727, 15: *B. finegoldii* DSMZ 17565, 16: *B. fragilis* BEI HM-20, 17: *B. fragilis* BEI HM-710, 18: *B. fragilis* BEI HM-711, 19: *B. fragilis* BEI HM-714, 20: *B. fragilis* NCTC 9343, 21: *B. intestinalis* DSMZ 17393, 22: *B. ovatus* ATCC 8483, 23: *B. ovatus* BEI HM-222, 24: *B. pectinophilus* ATCC 43243, 25: *B. plebeius* DSMZ 17135, 26: *B. salyersiae* BEI HM-725, 27: *B. sp.* BEI HM-18, 28: *B. sp.* BEI HM-189, 29: *B. sp*. BEI HM-19, 30: *B. sp.* BEI HM-22, 31: *B. sp.* BEI HM-23, 32: *B. sp.* BEI HM-258, 33: *B. sp.* BEI HM-27, 34: *B. sp.* BEI HM-28, 35: *B. sp.* BEI HM-58, 36: *B. stercoris* ATCC 43183, 37: *B. stercoris* BEI HM-1036, 38: *B. thetaiotaomicron* 3730, 39: *B. thetaiotaomicron* 3731, 40: *B. thetaiotaomicron* 633, 41: *B. thetaiotaomicron* 7330, 42: *B. thetaiotaomicron* 7853, 43: *B. thetaiotaomicron* 8702, 44: *B. thetaiotaomicron* 8713, 45: *B. thetaiotaomicron* 8736, 46: *B. thetaiotaomicron* 940, 47: *B. thetaiotaomicron* VPI 5482, 48: *B. thetaiotaomicron* wh302, 49: *B. thetaiotaomicron* wh305, 50: *B. uniformis* ATCC 8492, 51: *B. vulgatus* ATCC 8482, 52: *B. vulgatus* BEI HM-720, 53: *B. xylanisolvens* DSMZ 18836, 54: *P. distasonis* ATCC 8503, 55: *P. distasonis* BEI HM-169, 56: *P. johnsonii* BEI HM-731, 57: *P. johnsonii* DSMZ 18315, 58: *P. merdae* ATCC 43184, 59: *P. merdae* BEI HM-729, 60: *P. merdae* BEI HM-730.

**Extended Data Fig. 9.**
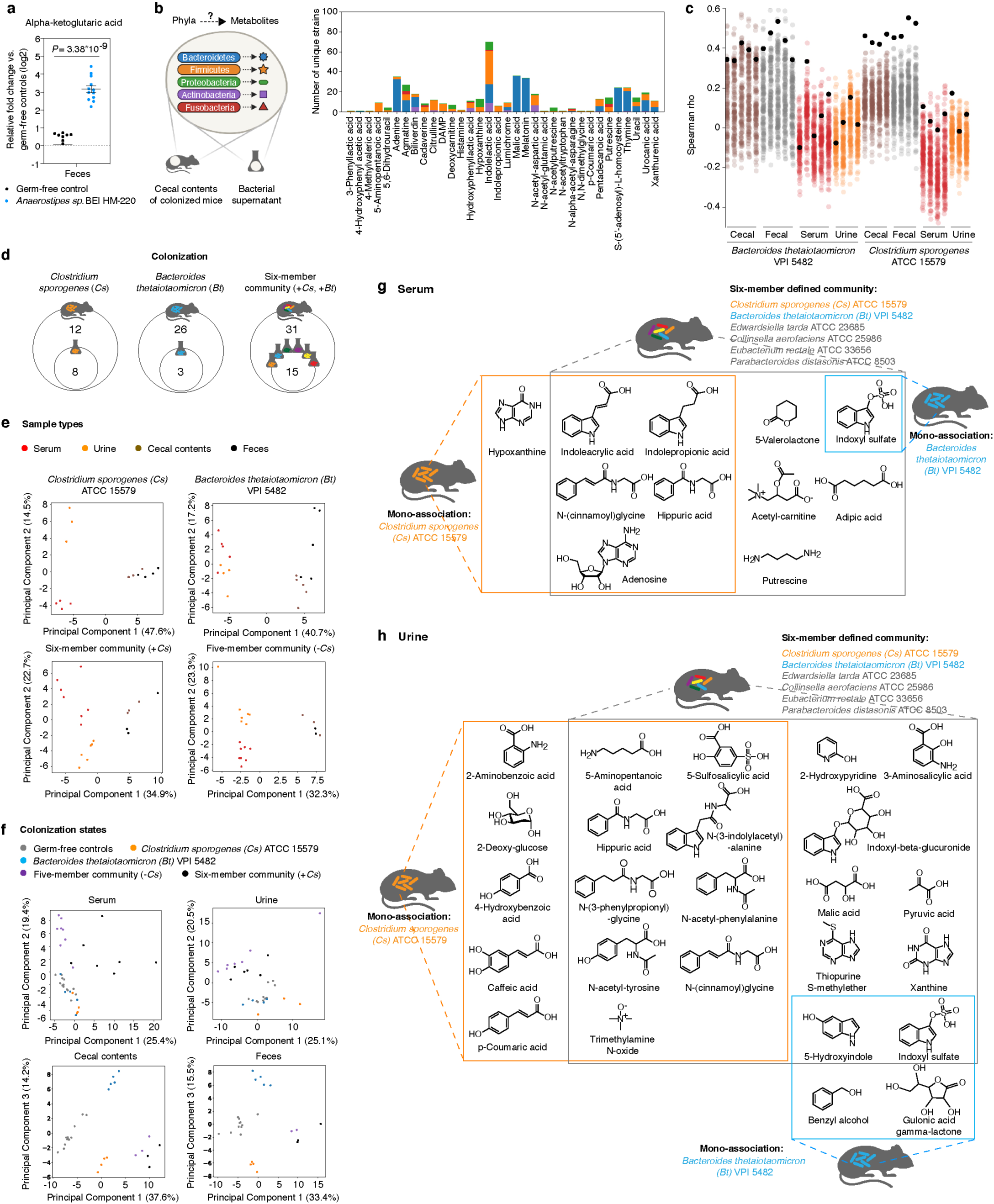
Metabolic contribution by individual gut microbes in a multi-species community. **a,** Alpha-ketoglutaric acid levels in feces of mice mono-colonized with *Anaerostipes sp.* BEI HM-220. Mean ± s.e.m. of two independent experiments, each with n = 4 mice (germ-free) or n = 5 or 7 mice (*Anaerostipes* mono-colonized). **b,** Left panel: MDMs were associated with specific bacterial phyla leveraging both *in vivo* and *in vitro* metabolomic data. Right panel: Number of mega-medium grown bacterial strains by phylum that produce MDMs identified in the cecal contents of mice colonized with *Bacteroides thetaiotaomicron* (*Bt*, n = 5), or *Clostridium sporogenes* (*Cs*, n = 3), or a six-member community (n = 3). Number of strains that produce at least one of these metabolites *in vitro* by phylum: Bacteroidetes: n = 52, Firmicutes: n = 60, Proteobacteria: n = 8, Actinobacteria: n = 16, and Fusobacteria: n = 3. Each metabolite shown was significantly produced both *in vitro* and *in vivo* (≥ 4-fold, corrected *P* < 0.05). Uniquely detected (non-coeluting) metabolites are shown (Supplementary Table 9). **c,** Spearman correlation between metabolomic profiles (standardized and scaled, log2-transformed, fold-change data) of individual *Bt-* or *Cs*-mono-associated host biofluids (cecal contents, feces, serum, or urine) and individual bacterial culture (158 mega-medium grown). Colored dots: Spearman’s rho values calculated by comparing metabolomic profiles of individual bacterial culture vs. individual biofluid of either *Bt-* or *Cs*-mono-associated mice. Black dots: Spearman’s rho calculated using metabolomic profiles of *Bt* or *Cs*, the same strains used for mono-association in mice. **d,** Venn diagram of overlapping metabolites that are significantly produced (≥ 4-fold, corrected *P* < 0.05) in culture and in the cecum of colonized mice. **e,** Principal component analysis (PCA) separates metabolomic profiles of identified metabolites by sample type in each colonization state. *P* values on metabolomic profile comparisons between different sample types of the same colonization state were determined using PERMANOVA: six-member community (*P* = 0.073), and all other colonization states (*P* = 0.001). **f,** PCA separates metabolomic profiles of identified metabolites by colonization states. *P* values on metabolomic profile comparisons between different colonization states of the same sample type were determined using PERMANOVA: *P* = 0.001 for all four sample types. **g**, **h**, Example chemical structures of significantly produced metabolites (≥ 4-fold, corrected *P* < 0.05) in serum (**g**) or urine (**h**) by each colonization state corresponding to Fig. 4b. *P* values in **a**, **b**, **d**, **g**, **h**: two-tailed *t*-test with Benjamini-Hochberg correction for multiple comparisons.

**Extended Data Fig. 10.**
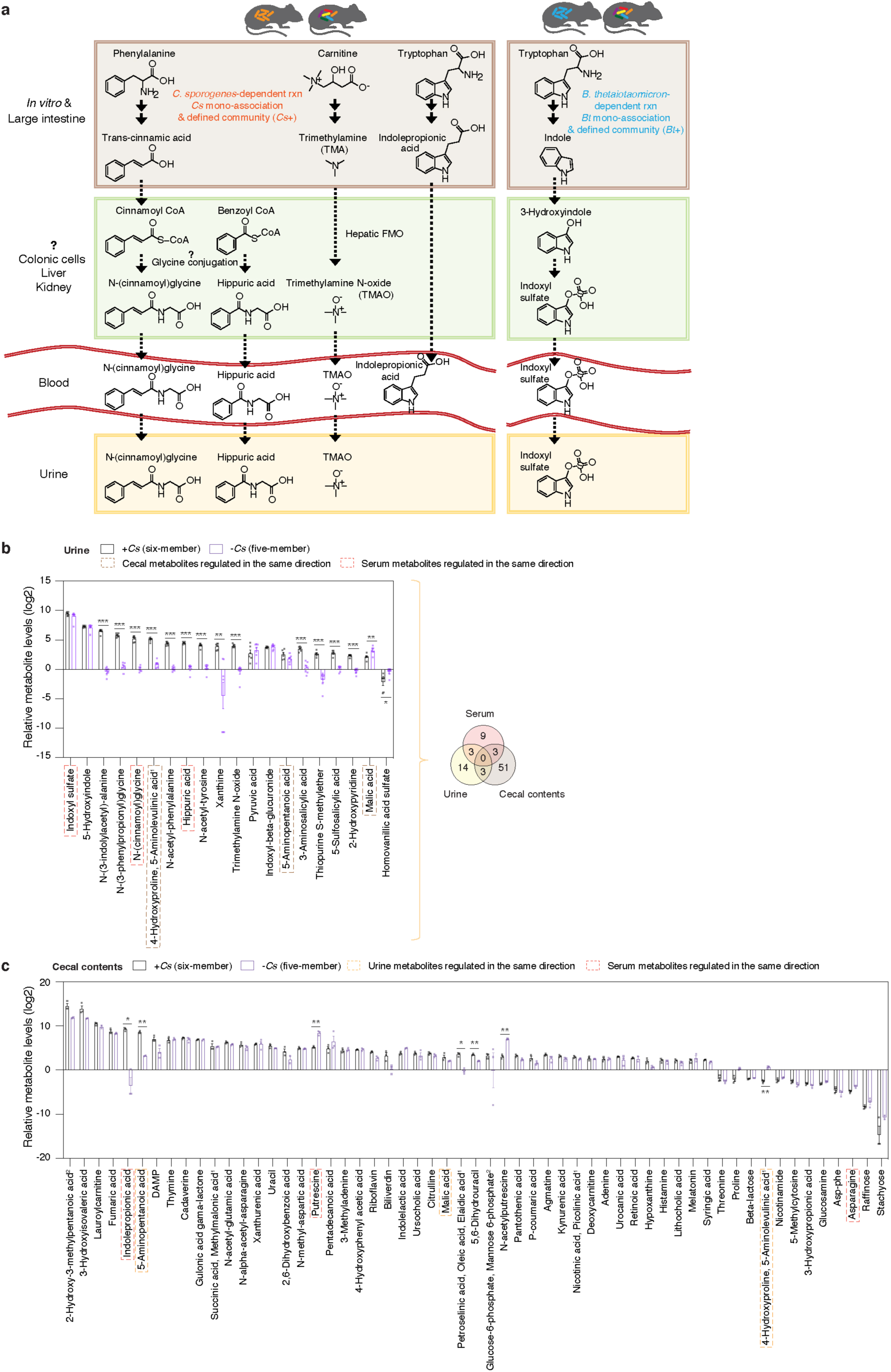
Metabolic contribution of multi-species communities in gnotobiotic mice. **a,** Proposed host-microbial co-metabolism pathways that could lead to the synthesis of specific host-microbial co-metabolites in the urine and serum of mice colonized with the six-member community. **b**, **c**, Metabolite levels in urine (**b**) and cecal contents (**c**) of mice colonized with the six-member community (+*Cs*) or the five-member community (-*Cs*). Metabolites shown represent a panel of significantly elevated or reduced metabolites (≥ 4-fold, corrected *P* < 0.05) in the six-member community. Superscript “1”: co-eluting metabolites as annotated in the mass spectrometry reference library (Supplementary Table 1). Superscript “2”: co-eluting isomeric metabolites with truncated names in the figure (“2-Hydroxy-3-methylpentanoic acid, 2-Hydroxy- 4-methylpentanoic acid”, and “Alpha-galactose 1-phosphate, Alpha-glucose 1-phosphate, Glucose-6-phosphate, Mannose 6-phosphate”). Mean ± s.e.m. of one experiment with n = 6 (urine, six-member community), n = 7 (urine, five-member community), and n = 3 (cecal, both six-member and five-member communities). **b**, **c**, *P* values: two-tailed *t*-test with Benjamini-Hochberg correction for multiple comparisons. * *P* < 0.05, ** *P* < 0.01, *** *P* < 0.001. Venn diagram (**b**) of significantly elevated and reduced metabolites in individual host biofluids (cecal contents, serum, and urine) using the same threshold in **b**.

**Supplementary Table 1.** Mass spectrometry compound m/z-RT reference library

|  |  |
| --- | --- |
| comp. info columns | dname, compound, peak, pubchem_CID, mz (precursor accurate mass/charge), rt (retention time identified on the qTOF instrument), adduct, molecular_formula, monoisotopic mass, QE_rt (retention time identified on the Q Exactive instrument), QE_ms2 (“*” denotes ms2 spectra generated on QE, spectra reported in Supplementary Table 3), qTOF_ms2 (“*” denotes ms2 spectra generated on qTOF, spectra reported in Supplementary Table 3) |
| comp. class. columns | kingdom, superclass, class, subclass |
| comp. id columns | canonical_smiles, inchikey |
| analytical_method | “mode”: C18 positive, C18 negative, HILIC positive |

**Supplementary Table 2.**
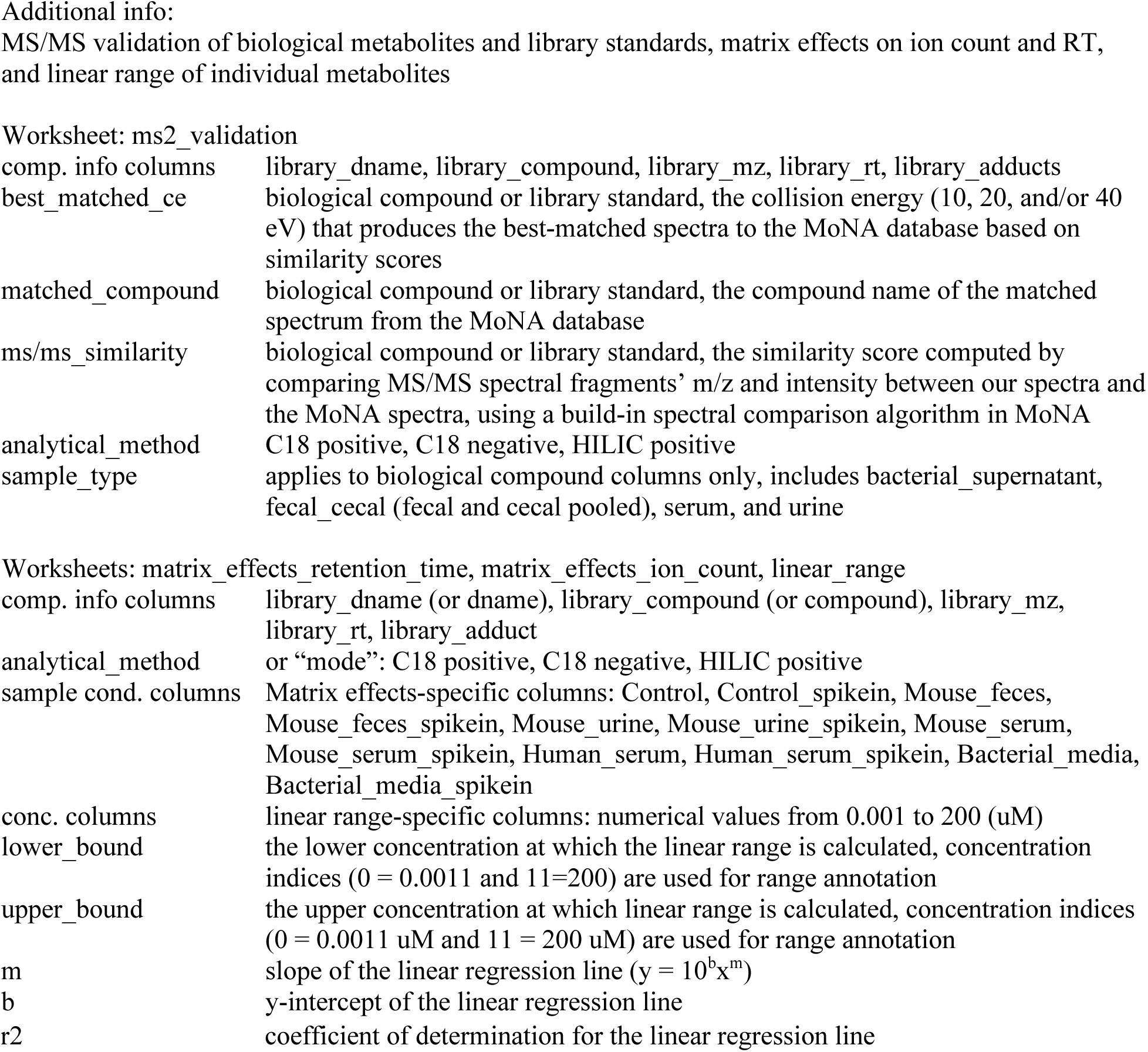

**Supplementary Table 3.** MS/MS spectra library constructed on qTOF and QE instruments

|  |  |
| --- | --- |
| Comp. info rows | Name, Precursor_mz, Precursor_type, Spectrum_type, InChIKey, SMILES, Formula |
| Instrument info rows | Instrument_type, Instrument, Ion_mode, Collision_energy |
| Spectra info rows | Num Peaks, m/z and intensity pairs separated by ',' |

**Supplementary Table 4.** Inter-instrumental retention time shift and correction

|  |  |
| --- | --- |
| comp info columns | library_dname, library_compound, library_mz, library_rt, library_adducts |
| corrected_rt | library rt corrected by polynomial transformation of the library based on inter-instrumental RT shifts of 10-20 robustly detected metabolites (e.g. internal standards) |
| measured_rt | RT measured on the second instrument |
| measured vs. library | measured RT on the second instrument minus library RT |
| measured vs. corrected | measured RT on the second instrument minus corrected library RT |
| analytical_method | C18 positive, C18 negative, HILIC positive |

**Supplementary Table 5.**
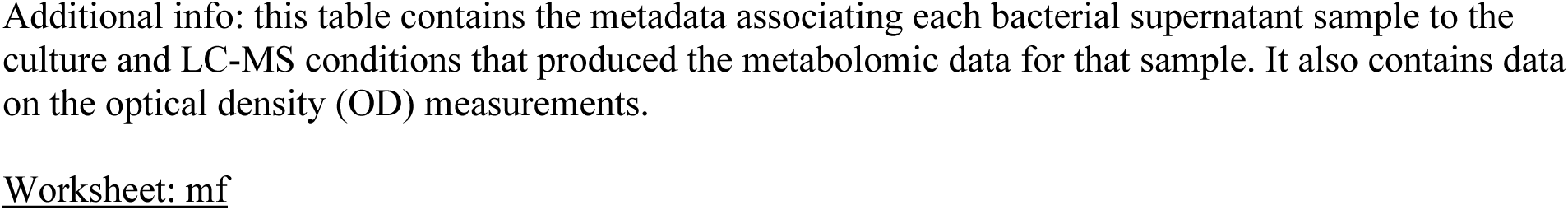
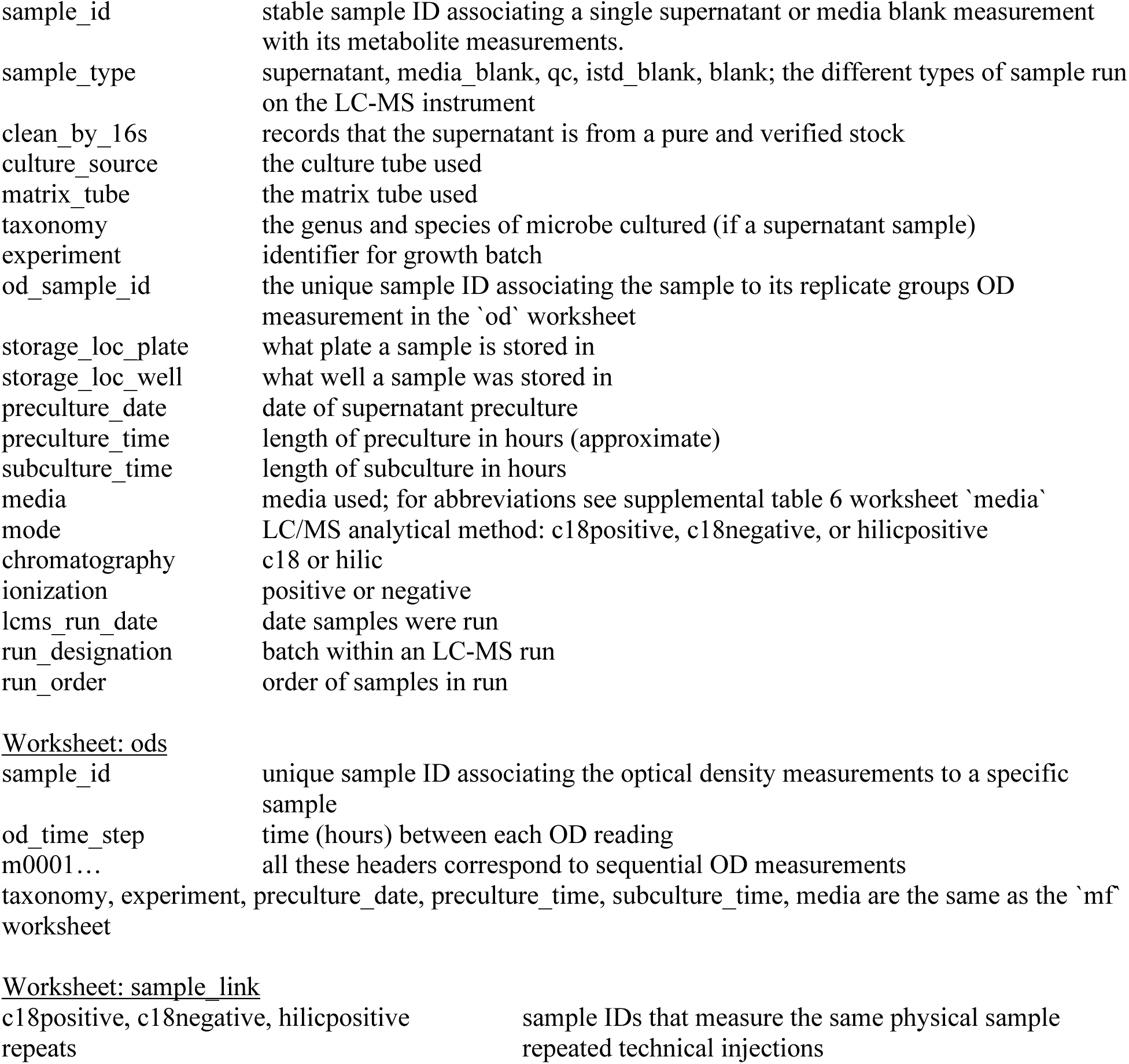
*in vitro* sample metadata

**Supplementary Table 6.**
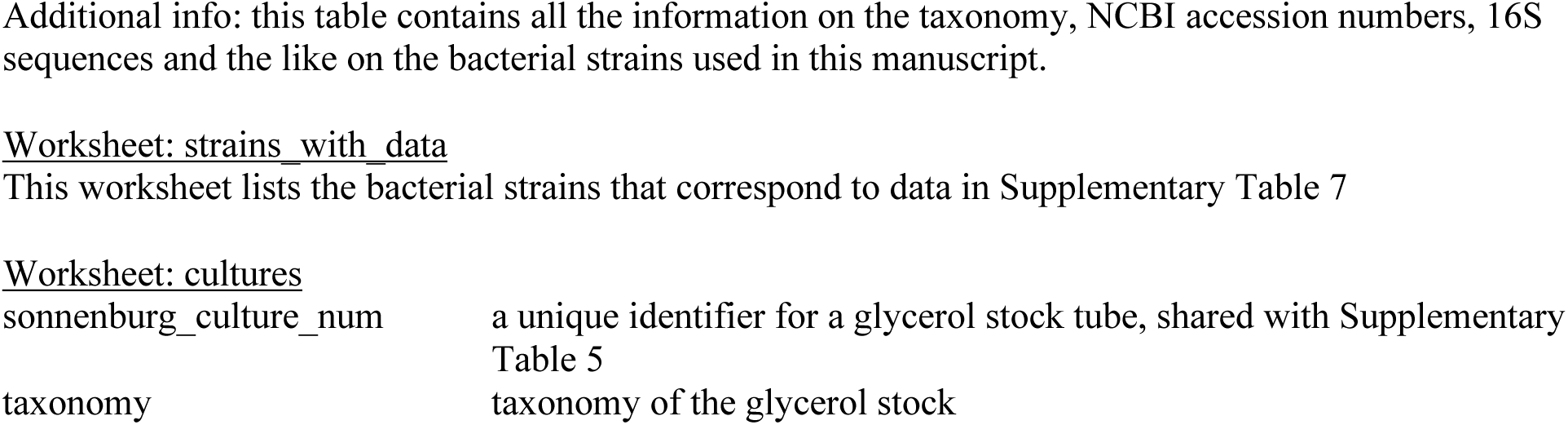
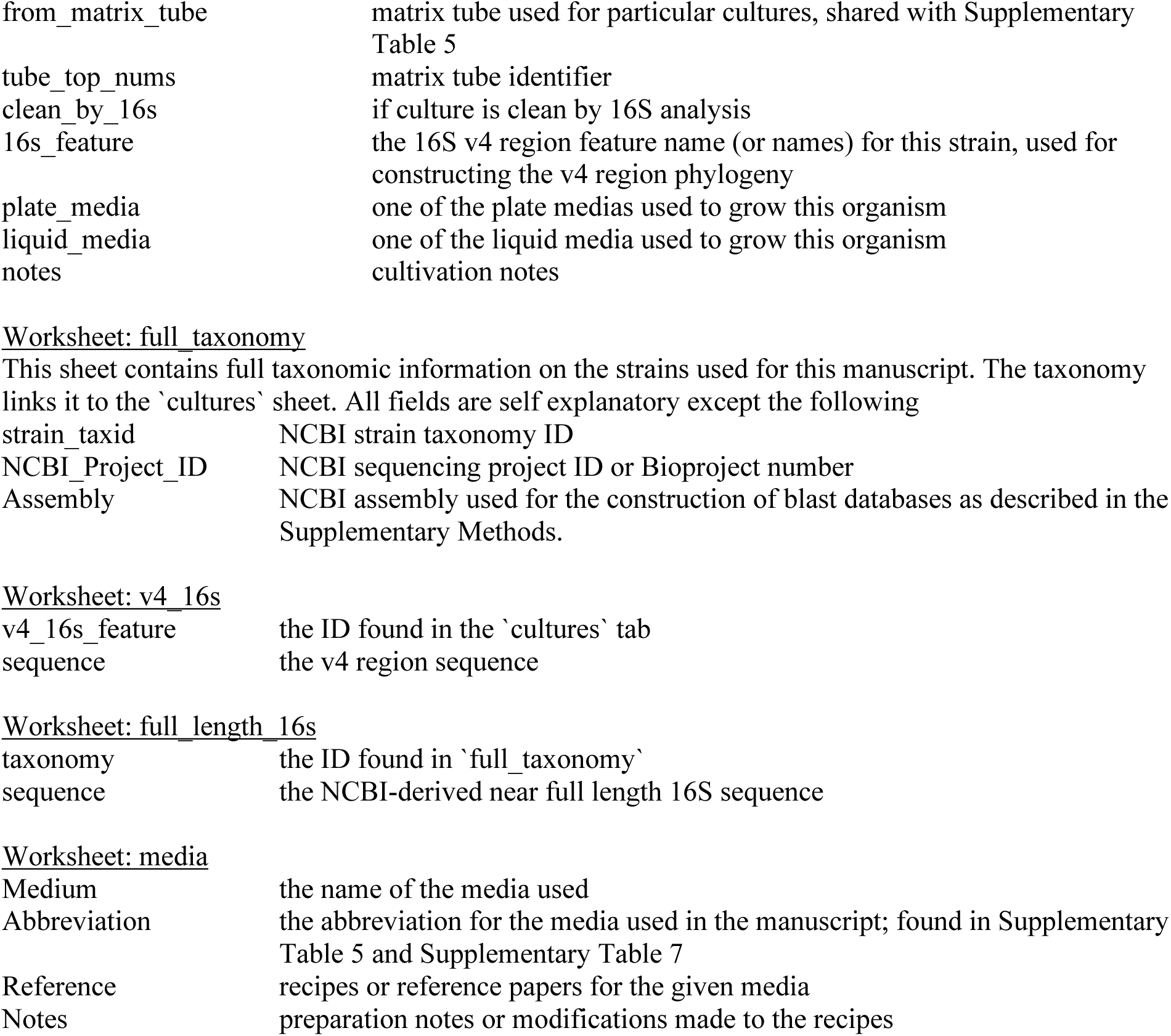
*in vitro* data matrices

Worksheet: cultures
|  |  |
| --- | --- |
| sonnenburg_culture_num | a unique identifier for a glycerol stock tube, shared with Supplementary Table 5 |
| taxonomy | taxonomy of the glycerol stock |
| from_matrix_tube | matrix tube used for particular cultures, shared with Supplementary Table 5 |
| tube_top_nums | matrix tube identifier |
| clean_by_16s | if culture is clean by 16S analysis |
| 16s_feature | the 16S v4 region feature name (or names) for this strain, used for constructing the v4 region phylogeny |
| plate_media | one of the plate medias used to grow this organism |
| liquid_media | one of the liquid media used to grow this organism |
| notes | cultivation notes |
| strain_taxid | NCBI strain taxonomy ID |
| NCBI_Project_ID | NCBI sequencing project ID or Bioproject number |
| Assembly | NCBI assembly used for the construction of blast databases as described in the Supplementary Methods. |
| Worksheet: v4\_16s |  |
| v4_16s_feature | the ID found in the 'cultures' tab |
| sequence | the v4 region sequence |
| Worksheet: full\_length\_16s |  |
| taxonomy | the ID found in 'full_taxonomy' |
| sequence | the NCBI-derived near full length 16S sequence |
| Medium | the name of the media used |
| Abbreviation | the abbreviation for the media used in the manuscript; found in Supplementary Table 5 and Supplementary Table 7 |
| Reference | recipes or reference papers for the given media |
| Notes | preparation notes or modifications made to the recipes |

**Supplementary Table 7.**
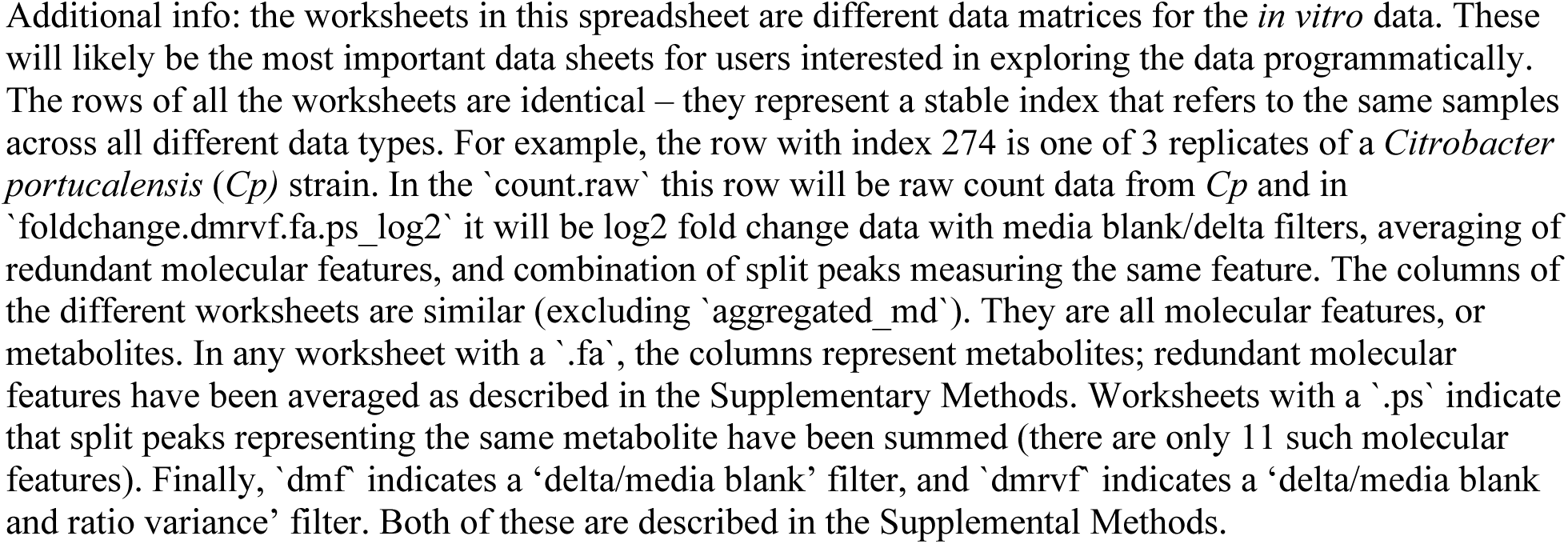
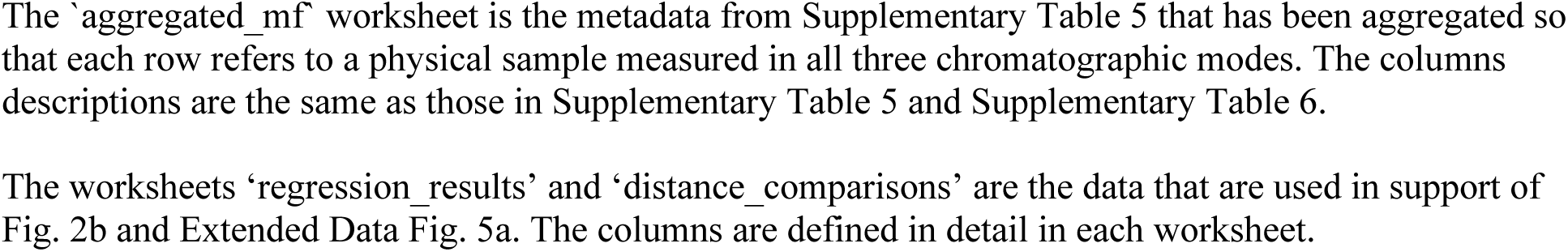
*in vitro* data matrices

**Supplementary Table 8.** The *in vivo* database and *in vivo* pipeline output data matrices

|  |  |
| --- | --- |
| sample_id | assigned to the physical sample |
| run_id | assigned to the physical sample analyzed with different methods |
| ms_dial_sample_name | names used in the msdial analysis output file |
| method columns | chromatography, ionization |
| sample_type | serum, urine, caecal, feces |
| colonization | germ-free, Bt, Cs, Bt_Ca_Er_Pd_Et, Cs_Bt_Ca_Er_Pd_Et, conventional, Cp, As |
| experiment | mono-colonization, community, conventional, mono-colonization_2 |
| collection_time | denotes independent experimental repeats |
| mouse_id | community database-specific column, denoting individual mouse |
| tissue_measurement | community database-specific column, denoting three sections of the cecum |
| c18positive | metadata-specific column, run ids associated with c18positive method |
| c18negative | metadata-specific column, run ids associated with c18negative method |
| hilicpositive | metadata-specific column, run ids associated with hilicpositive method |
| metabolite columns | fc_matrix-specific: fold-change (log2-transformed) values of individual metabolite of colonized vs. germ-free controls, specific to each sample type and analytical method, one column per metabolite per analytical method, in dnames<br>istd_corr_ion_count_matrix-specific: istd-corrected raw ion counts of individual metabolite for all sample types and colonizations states, one column per metabolite per analytical method, in dnames<br>mode_collapsed_fc_matrix: fold-change (log2-transformed) values as described in the fc_matrix above, but with one column per metabolite. For individual metabolites detected by multiple analytical methods, their fold-change values were averaged among the preferred detection methods (See Supplementary Method for details). Column names are compound names, with co-eluting compounds separated by commas |

Worksheets:
mouse sample databases: db\_mono-colonization, db\_community, db\_conventional
*in vivo* sample metadata: metadata
*in vivo* pipeline output data matrices: raw\_ion\_count\_matrix (columns are dnames, prior to averaging cecal section data in community experiment), istd\_corr\_ion\_count\_matrix (columns are dnames), fc\_matrix (fold change, log2-transformed, in dnames), mode\_collapsed\_fc\_matrix (fold change, log-2 transformed, columns are compound names)

**Supplementary Table 9.**
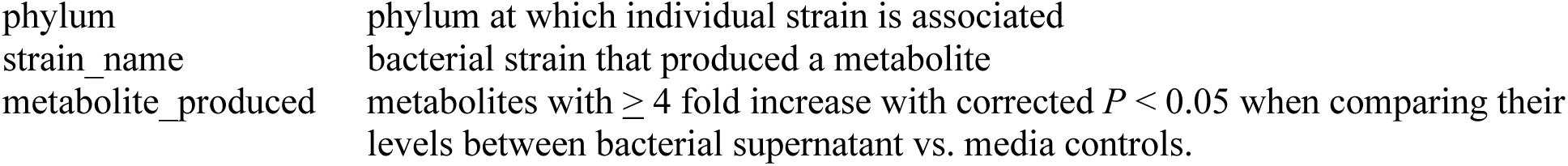
List of “phylum-associated” metabolites

|  |  |
| --- | --- |
| phylum | phylum at which individual strain is associated |
| strain_name | bacterial strain that produced a metabolite |
| metabolite_produced | metabolites with $\geq 4$ fold increase with corrected $P < 0.05$ when comparing their levels between bacterial supernatant vs. media controls. |

## Supplementary Methods

Title: Supplementary Methods

1. Mass spectrometry LC/MS methods

○ Instrumental and chromatographic settings
○ Metabolomics sample preparation
2. Data analysis

○ Custom bioinformatics: *in vitro* pipeline
○ Custom bioinformatics: *in vivo* pipeline
3. Bacterial sequencing and phylogenetics

○ Purity analysis
○ Phylogenetics
4. Distance comparisons and classifiers

○ Distance calculations
○ Correlation between labels in phylogenetic and metabolomic trees
○ Distance comparisons
○ Random forest classifications
5. Mega Medium (MM) preparation protocol
6. Salyer’s Minimal Medium (SMM) preparation protocol

