## Supplementary Methods for "A metabolomics pipeline enables mechanistic interrogation of the gut microbiome"

#### Mass spectrometry LC/MS methods

*Instrumental and chromatographic settings.*

Published C18 methods[1] and HILIC method[2] were implemented with minor modifications. The C18 positive method (ESI+) used mobile phase solvents (LC-MS grade) consisting of 0.1% formic acid (Fisher) in water (A) and 0.1% formic acid in methanol (B). The C18 negative method (ESI-) used mobile phase solvents consisting of 6.5 mM ammonium bicarbonate (Sigma) in water at pH 8 (A) and 6.5 mM ammonium bicarbonate in 95:5 v/v methanol:water (B). For both C18 methods, the gradient profile was from 0.5% B to 70% B in 4 minutes, from 70% B to 98% B in 0.5 minutes, and holding at 98% B for 0.9 minute before returning to 0.5% B in 0.2 minutes. The HILIC (Hydrophilic Interaction Liquid Chromatography) positive method (ESI+) used mobile phase solvents consisting of 0.125% formic acid and 10 mM ammonium formate (Sigma) in water at pH 3 (A) and 0.125% formic acid in 10 mM ammonium formate in 95:5 v/v acetonitrile:water (B). The gradient profile was held at 100% B for 2 minutes, from 100% B to 70% B in 5 minutes, holding at 70% B for 0.7 minute, from 70% B to 40% B for 1.3 minutes, holding at 40% B for 0.5 minutes, from 40% B to 30% B for 0.75 minutes, before returning to 100% B for 2.5 minutes and holding at 100% B for 4 minutes. The flow rate was 350 µL per minute (C18 positive, C18 negative), and 400 µL per minute (HILIC positive). The sample injection volume was 5 µL (C18 positive, C18 negative), and 3 µL (HILIC positive). LC separations were made at 40$^{o}$C on separate columns fitted with a Vanguard pre-column of the same composition dedicated to each analytical method: Waters Acquity BEH 1.7 µm particle size, 2.1 mm id x 100 mm length (C18), Waters Acquity BEH Amide 1.7 µm particle size, 2.1 mm id x 150 mm length (HILIC). For all analytical methods, data were collected at a mass range of 70-1000 m/z at an acquisition rate of 2 spectra per second. Specific ion source parameters included Fragmentor (140V), Gas Temp (250$^{o}$C), Sheath Gas Temp (200$^{o}$C), and VCap (4000V).

*Metabolomics sample preparation.*

All samples were stored at -80$^{o}$C until use, and were thawed on ice immediately before extraction. 200 µL of serum and bacterial supernatant samples were used for extraction directly without dilution. Urine samples were diluted 1:20 in LC-MS grade water (Fisher) to reach a final volume of 200 µL prior to extraction. Protein precipitation for bacterial, serum, and urine samples was conducted by adding 1 mL of extraction buffer (see composition below) in 100% methanol (LC-MS grade, Fisher) to 200 µL of each sample in a 2 mL 96-well microplate (Fisher), sealed with a silicone mat (Agilent), and vortexed to mix. For feces and cecal contents, ~ 20 mg of feces or cecal contents were added to ~ 20 mg of acid-washed glass beads (150-212 µm, Sigma) in a 2 mL autoclaved screw top vial. In the same vial, 600 µL of water (LC-MS grade, Fisher) and 600 µL of recovery buffer in 100% methanol (see composition below) were added. Fecal and cecal slurries were homogenized at 4$^{o}$C using a Mini Beadbeater operating at 3,500 oscillations per minute for 5 minutes. For all sample types, samples were subsequently incubated at room temperature for 5 minutes, followed by centrifugation at 5,000 x g for 10 minutes. Two 440 µL aliquots of the same supernatant were transferred to two separate 2 mL plates and dried under air in a Biotage TurboVap. One of these dried plates was sealed and archived at -80$^{o}$C. The dried extracts were reconstituted in 200 µL reconstitution buffer (see composition below) in 50% methanol in water (v/v, LC-MS grade, Fisher) by vortexing at max speed for 5 seconds. Reconstituted sample extracts were centrifuged at 2,000 x g for 1 minute, and filtered through a 96-well Durapore PVDF 0.22-um filter plate (Millipore) into in a 1 mL 96-well plate (Agilent) by centrifugation at 2,000 x g for 10 minutes. Plates were then sealed with 96-well cap mats (Agilent) and stored at -80ºC until LC-MS analysis. QC samples were generated by pooling 4 µL from each well of the experiment into a single designated well on the same plate for LC-MS analysis.

The extraction buffer consisted of 4-Chloro-phenylalanine (6.8 µM, Carbosynth), Tridecanoic acid (6.8 µM, Sigma), and 2-Flurophenylglycine (3.4 µM, SCBT) in 100% methanol. The reconstitution buffer included the internal standards: Phenylalanine-2,3,4,5,6-d5 (12.5 µM, CIL), Glucose-1,2,3,4,5,6,6-d7 (25 µM, CIL), Methionine-methyl-d3 (12.5 µM, CIL), 4-Hydroxyphenyl-d4-alanine (3.125 µM, CDN), Tryptophan-2,4,5,6,7-d5 (12.5 µM, CDN), Leucine-5,5,5-d3 (12.5 µM, CDN), N-Benzoyl-d5-glycine (6.25 µM, CDN), 4-Bromo-phenylalanine (12.5 µM, Sigma), Progesterone-d9 (3.125 µM, CIL), Di-N-octyl phthalate-3,4,5,6-d4 (12.5 µM, CDN), d19-Decanoic acid (12.5 µM, CDN), d15-Octanoic acid (25 µM, CDN), Indole-2,4,5,6,7-d5-3-acetic acid (12.5 µM, CDN), Carnitine-trimethyl-d9 (3.125 µM, CDN), and d27-Tetradecanoic acid (12.5 µM, CDN), in 50% methanol in water. The final concentration for each internal standard in these buffers was determined by choosing a concentration falling within its linear dynamic range as measured by each analytical method.

**Data analysis**

*Custom bioinformatics: in vitro pipeline*

The *in vitro* metabolomics pipeline was designed to enable comparison across experimental and analytical batches. Five main quality control steps were used (in order): removal of samples with high intra-replicate variability, removal of samples with substantial intra-experimental variation, internal standard and media blank selection and normalization, selection of measured molecular features (metabolites) on the basis of intra-replicate and intra-experiment variability, and removal of samples based on RF classification accuracy of taxonomy. Code is available in 'in_vitro_pipeline.ipynb', depicted in Extended Data Fig. 2, and summarized below.

Individual replicate groups were compared by Pearson's correlation coefficient. Intra-replicate ion count (area under the curve) variability was generally low, with average correlation coefficient of 0.9-1.0 (log transformed data) depending on batch (Extended Data Fig 4b). Individual replicates were removed if their correlation with the other replicates was two standard deviations below the mean of all replicates. Multiple thresholds and removal strategies produced similar results. Entire batches (a set of samples that were harvested and ran on the instrument together) were then compared to identify outlier samples. Samples were compared via Principal Component Analysis (PCA). Samples distant from the centroid of the first two PCs were inspected at the chromatogram level, with focus on metabolites with high loadings on those PCs. In general, this strategy identified samples with substantial differences in single metabolites due to technical artifacts (e.g. poor peak integration, non-detection, column pressure fluctuation, etc.).

LC-MS measurements are frequently confounded by matrix effects (differences in ionization efficiency for the same metabolite present in different biological sample types) and changes in instrumental sensitivity over time. To minimize these effects, we normalized samples based on a shared set of internal standards (IS). After sample removal, IS ion counts were compared within and between experiments on a per-analytical method basis. Based on matrixeffect experiments, we assumed that highly variable (>10X difference in count between samples, intra-experiment) IS were not representative of ionization efficiency and were excluded from analysis. The remaining panel of IS (2-Flurophenylglycine, 4-Bromo-phenylalanine, 4-Chloro-phenylalanine, d19-Decanoic acid, Methionine-methyl-d3, N-Benzoyl-d5-glycine, Indole-2,4,5,6,7-d5-3-acetic acid, Phenylalanine-2,3,4,5,6-d5, and Tridecanoic acid) were summed (per sample) and used to compute a multiplicative scaling factor to equalize each sample ($\frac{\overline{\sum IS}}{\sum_{sample} IS}$). This was done on a per-analytical method basis. For example, the correction factor for C18 positive method was computed from only IS detected in C18 positive method and applied to only detected features using this method. Counts of all metabolites (per sample and per method) were multiplied by the samples (method-specific) scaling factor prior to further analysis. We explored other normalization strategies including a scaling factor based on a weighted sum of IS, but chose total IS sum because it minimized intra-replicate coefficients of variation (CV).

In addition to IS normalization, we computed several other data transformations with the goal of allowing inter-experiment comparisons (Supplemental Table 7). IS-based correction helped to normalize average molecular feature intensity across runs but could not correct for all inter-experiment variation or biological differences, e.g., differences in molecular feature count due to media batch. For *in vitro* data used in the figures (e.g., Fig. 3) and in the Metabolomics Data Explorer, the data were subjected to the following transformation to mitigate these concerns. First, a 'delta' matrix was constructed (IS-corrected supernatant samples minus IS-corrected media blanks on a per-molecular feature basis). For each replicate group (i.e. three biological replicates), molecular feature counts were eliminated if the replicate group average was less than 3000 counts or the delta was less than 3000 counts (absolute value). The purpose of this filter was to eliminate peak counts that were likely artifactual (e.g., due to integration of a non-peak or a peak with low signal-to-noise ratio. This filter eliminated 9.9% of measured values. Different filters (1000-10000 count) were explored with minimal impact on the data since almost all of these counts were random noise or very low quality peaks. Second, molecular features that varied by more than 10X (by ion count) within a replicate group were eliminated. This threshold was chosen as the elbow of the distribution of (per feature) per replicate group maximum count divided by minimum count. This eliminated 1.0% of the measured values. Finally, we used this filtered data and computed a fold change matrix. Values in the fold change matrix are the count of a molecular feature in a supernatant sample divided by the count of that feature in the blank media that was used to grow that supernatant. This final matrix enables inter-experimental comparisons while minimizing technical and biological batch effects. All intermediate transformations and the raw untransformed data are available in Supplementary Table 7.

Because many metabolites were detected by more than one analytical method, we next identified a strategy for selecting which method(s) to prefer for fold-change data on a per-metabolite basis. In this step, columns of the data matrix were transformed from detected molecular features detected in multiple methods (redundant) to a single metabolites (non-redundant). For each molecular feature detecting the same metabolite we computed the CV (calculated on a per-replicate basis) for all replicates with those features. We compared the distributions of CVs and the linear correlation between molecular features. In cases where correlation was low, (Pearson r < ~0.4) we selected the feature with the better CV distribution (smaller mean, positively skewed) and ignored the other molecular feature. In cases where correlation was high, we averaged the fold changes. When a metabolite was detected by only one molecular feature it was included regardless of how well that feature was correlated with the other detecting features.

The final quality control step evaluated the similarity of supernatant samples to all other samples from the same phylum. Preliminary work with our data showed that phylum level differences were evident in our supernatants, and they could be easily identified by random forest classifiers (RFs). Using a test:training split of 1:2, RF classifiers were trained to predict the phylum of the microbe generating each supernatant sample (see 'sample_qc_classification.ipynb'). The phylum of each test sample was then predicted and misclassifications inspected. This analysis was repeated with different parameters (e.g., binary classification or multiclass classification) and with different input data (e.g., fold-change or ion count data). Consistently, misclassified samples were individually inspected for phyla-specific features (e.g., samples were contaminated by Indolepropionic acid-producing *C. sporogenes* from Phylum Firmicutes). Samples with significant counts (>10,000 count average across all replicates) of phyla-specific metabolites (from a different phyla) that did not replicate on repeated growth were eliminated.

Here we have presented this methodology as linear, but it should be noted that the filtration process was iterative with later steps informing earlier filtration steps. As an example, the PCA and RF analysis steps revealed molecular features that were inconsistent across runs and had to be eliminated in the initial data curation. In total, this pipeline represents a conservative approach more likely to eliminate 'good' samples than allow samples containing substantial artifacts (biological or technical) to remain in the data. Cumulatively this pipeline eliminated 8.13% of samples and 10.9% of measured counts in the retained samples.

*Custom bioinformatics: in vivo pipeline.*
The *in vivo* pipeline integrated data across mono-colonization, community, and conventional mouse experiments and three analytical methods to generate a unified, metabolite fold-change matrix. First, the sample database and MS-DIAL output files from the experiments were parsed and integrated into a single data matrix of raw ion counts across the three analytic methods. Based on the reference library, a small subset of metabolites exhibited dual peaks specific to an analytical method. For these metabolites, the raw ion counts from these two peaks were summed in the raw ion count matrix. Next, a shared set of internal standards were used to normalize across experiments to account for inter-experimental variations in instrument sensitivity. As the qTOF instrument was in a shared core facility that ran near full capacity, not a dedicated instrument for these metabolomics methods, run-to-run variation in sensitivity was an important issue. To account for these variations, the raw ion count from each metabolite was normalized to the sum of internal standard ion counts specific to each experiment (e.g., community experiment) and sample type (e.g., serum). The fold change matrix was generated next by calculating the relative fold change between metabolite ion counts detected in colonized mice vs. germ-free controls for each experiment and sample type. In particular, cecal samples from the community experiment were processed differently from other experiments, because the cecum from each mouse was trisected and three cecal samples were harvested per mouse. In this case, before calculating fold-change values, the three cecal samples for each mouse were collapsed into a single sample row in the ion count matrix by averaging the ion counts for each metabolite. Lastly, when a metabolite was detected by multiple analytical methods, its fold-change values were averaged among the preferred detection methods, which were determined based on the consistency (CVs) of detection among replicates for each method (see *in vitro* pipeline).

The final fold-change matrix was used for several analyses: 1) PCA analyses to separate metabolomic profiles by sample types or colonization states, 2) statistical calculation to identify significantly regulated metabolites (≥ 4 fold) using *t*-test with Benjamin-Hochberg corrections for multiple comparisons ($P < 0.05$), and 3) violin plots to visualize metabolites associated with a specific colonization state in a given sample type. All steps detailed here are explained in depth in the Jupyter notebook "mouse_data_analysis.ipynb". The final fold-change matrix was used to construct Fig. 4 and Extended Data Figs. 8e, 9, 10.

Fold-change matrices generated by both *in vitro* and *in vivo* pipelines can be interactively accessed in the web-based Metabolomics Data Explorer (https://sonnenburglab.github.io/Metabolomics_Data_Explorer)

**Bacterial sequencing and phylogenetics**

*Purity analysis*
Culture purity was assessed with 16S amplicon sequencing of the V4 (515f, 806r) region using Golay barcoded primers following the protocols of the Earth Microbiome Project (https://earthmicrobiome.org/). Briefly, DNA from bacterial cultures was extracted using DNeasy PowerSoil HTP 96 Kit (Qiagen). PCR conditions were as follows: initial denaturation 3 minutes at 94°C, followed by 35 cycles of denaturation (45 seconds at 94°C), annealing (60 seconds at 50°C), extension (90 seconds at 72°C), followed by a final elongation step of 10 minutes at 72°C. PCR products were cleaned using UltraClean 96 PCR Cleanup Kit (Qiagen) and quantified with the Quant-iT PicoGreen dsDNA Assay Kit (Invitrogen). Cleaned products were pooled in equimolar amounts and sequenced using 300-bp paired end reads on a MiSeq (Illumina). Illumina reads were demultiplexed and sequence variants were determined using QIIME 1.9.1[3] and DADA2[4]. Sequence variants in each sample were compared against the expected sequence of the grown strain taken from NCBI. A sequence match was considered to be at least 251 nucleotides (nt) out of 253 nts from the NCBI reference. If more than 5/1000 reads did not match, the culture was considered contaminated. Some exceptions for nt match thresholds were made for strains with low quality 16S sequences in the NCBI database (e.g. *Tyzzerella nexilis*).

*Phylogenetics*
Phylogenies were constructed with both V4 amplicons generated by Illumina sequencing (above) and with near full-length 16S sequences identified via automated and manual search of NCBI. 16S sequences >1200 nts in length were considered for full-length phylogeny reconstruction and strains were excluded if a full-length sequence could not be found. In both cases, phylogeny was constructed using Clustal Omega with default parameters using the EMBL-EBI web interface[5].

Because strains frequently had more than one associated V4 16S variant, we created a modified phylogenetic tree with a one-to-one correspondence between tips and strains used in this study that was used in the distance calculations described below (see 'phylogenetic_tree_modifications.ipynb'). Trees were visualized with iTOL[6].

**Distance comparisons and classifiers**

*Distance calculations*

To generate distance matrices used for Fig. 2a, and Extended Data Fig. 7a, d-i, we followed the following procedure (detailed in 'distance_comparisons.ipynb'). First, we averaged the measured values for metabolites for each strain across all experiments and log transformed the data. For each pair of strains, we calculated the Euclidean distance between the metabolite vector of those strains, excluding metabolites that were not measured in both (e.g., metabolites with NaNs in one or both vectors were removed prior to calculation). We also computed the tip-to-tip distances for all strains using the V4 region and full-length phylogenetic trees using scikit-bio. We also computed metabolomic distance matrices using input metabolomic data with various transformations and levels of filtration. Specifically, we used count data, unfiltered fold change data, and fold change data with media blank, delta, and variance ratio filtration. The outcome of the following analyses was unaltered with these different input data sets suggesting our results are robust and not the product of a well-chosen transformation.

The heatmap in Extended Data Fig. 6a with 158 mega-medium grown strains hierarchically clustered based on their metabolomic distance was generated as described in 'in_vitro_heatmap_scatter_plot.ipynb'. The *in vitro* metabolite fold-change matrix (log2 transformed) was used. On the y-axis, taxonomies, or bacterial strains, are clustered by their metabolomic distance. We first computed the metabolomic distance matrix using the Euclidean distance between fold-change values as the distance metric. We then hierarchically-clustered (Ward's method) the taxonomies and constructed the heatmap according to the resulting taxonomy order. On the x-axis, metabolites are also hierarchically-clustered (Ward's method) using the Euclidean distance between the fold-change (log2 transformed) values across all taxonomies.

Generation of metabolomic trees weighted based on chemical similarity between the metabolites in the *in vitro* dataset (Extended Data Fig. 7b, c) used the procedure as described in 'invitro_metabolite_similarity.ipynb'. We first computed a chemical distance matrix using pairwise Tanimoto 2D structural similarity scores (PubChem database scores range from 0 to 100 as the most similar) among all uniquely detected (non-coeluting) metabolites in the *in vitro* dataset. Based on this chemical distance matrix, we performed hierarchical clustering on the metabolites and assigned weights to each metabolite such that the sum of weights within each cluster of closely related metabolites equal 1. Lastly, we constructed weighted metabolomic trees using metabolomic profiles of bacterial taxa that are hierarchically-clustered based on their weighted Euclidean metabolomic distance.

To compare the weighted and unweighted metabolomic trees, we performed the Mantel test to correlate the two distance matrices computed on the weighted and unweighted Euclidean distance. We found that the matrices were highly correlated despite the addition of weighting. To transfer the metabolomic trees to iTOL for visualization, we converted the metabolomic clustering (expressed as a linkage matrix) into Newick trees, which iTOL supports as input. Supporting code is described in 'invitro_metabolite_similarity.ipynb'.

*Correlation between labels in phylogenetic and metabolomic tree*

To understand if the relative position of the labels (strains and associated taxonomies) were similar between the phylogenetic and metabolomic tree we used the following procedure (Extended Data Fig 7a). For each tree we calculated the 4th percentile of the tip-to-tip distance (TTD) matrix. We designated this value as the 'radius' of a 'neighborhood' such that for a given tip, any other tip with pairwise distance less than the radius was considered part of the given tips neighborhood. For each tip, we compared the neighborhoods of that tip in the phylogenetic and metabolomic tree. We calculated the overlap between these neighborhood sets at each taxonomic level. We used the number of tips in the phylogenetic neighborhood as the denominator to calculate the fraction of the labels seen in the metabolomic neighborhood that were found in the phylogenetic neighborhood. For example, if the phylogenetic neighborhood contained 5 Bacteroidetes and the metabolomic neighborhood 10 Bacteroidetes, the fractional representation would be 1.0 at the phylum level. If the phylogenetic neighborhood contained *B. ovatus* strain 1, *B. ovatus* strain 2, and *B. xylanisolvens* strain 1 and the metabolomic neighborhood contained *B. ovatus* strain 1, *B. ovatus* strain 10, and *P. distasonis strain* 1, the overlap would be 1/3 at the strain level and 2/3 at the species level. We repeated this procedure for all tips in the tree, and for definitions of neighborhood radius ranging from the 1st to 10th percentile of distances (in 1% increments). The results shown in Extended Data Fig. 7a are representative and are from a radius equal to the 4th percentile in each TTD respectively.

To calculate how significant the observed fractional coverage was, we used a permutative approach. For each tip, we calculated the true fractional overlap between neighborhoods. We then shuffled the labels of the TTD 1000 times, and recorded the number of times a fractional overlap larger than the observed occurred. The p-value (shuffled fractional overlap > observed fractional overlap / 1000) is reported in Extended Data Fig. 7a. The code for this section can be found in the github repository under "distance_comparisons.ipynb".

*Distance comparisons*

For Extended Data Fig. 7a, d-i we compared the phylogenetic and metabolomic distances. Patterns were consistent across multiple different transformations of the data including data type (fold change or count), data filtration (no filter or variance and intensity filter), and NaN removal (no removal or replacement by 1). The LOESS regressions of the data matrices were done using scipy.

*Random forest classification*

To establish if there were consistent metabolomic signals of taxonomic labels (e.g., Bacteroidetes or Firmicutes) we used random forest (RF) classifiers. For Fig. 3a, 25 RFs were trained on each taxonomic level (e.g. to predict phylum labels) using metabolomic data (see 'figure_3_classifiers.ipynb'). Fold change data were used to avoid batch effects of raw ion counts, though classification accuracy was not significantly different using count data. Data were split 1:2 test:training without class balancing due to the over-representation of Bacteroidetes and Firmicutes in the data.

For identification of Bacteroidetes specific metabolites, we trained RFs to predict phylum level labels of Bacteroidetes and Other (binary classification, Extended Data Fig. 8a, top panel). This strategy identified Bacteroidetes specific metabolic behaviors with high accuracy leading to ~97% classification accuracy on average. We built 50 forests and averaged the feature importance from all forests to produce the overall importance data in Extended Data Fig. 8a, bottom panel.

To avoid overfitting both of the above tasks, we significantly constrained the number of total decisions a tree could make (maximum depth ≤ 5) and eliminated features that were present in less than 30% of samples. Other thresholds yielded similar results, though below this depth, trees lost accuracy.

**Mega Medium (MM) preparation protocol**

This recipe was adapted from a previous publication[7] and from a personal communication of N. McNulty. The starch was used as indicated in the supplemental methods.

Steps

1. Add dry ingredients, liquid ingredients, and mix well (table 1).
2. pH to ~7.0. In practice, this takes slightly less than 2.5 mL 10M KOH per 500 mL media. After pH measurement, add 3.6 – X mL water where X is mL KOH added.
3. If plates are needed, add 7.5 g agar per 500 mL of media.
4. Autoclave cycle 4 (250 F for 25 minutes at 20 PSI, 20 minutes dry).
5. As soon as autoclaving starts, remove vitamin mix from freezer to let thaw. Once thawed, make vitamin solution and filter sterilize (table 2).
6. After media has cooled, sterilely add vitamin solution.

**Table 1**

| Component | Amount (in 500 mL) | Final conc. | Mfr. | Vendor (cat. #) |
|---|---|---|---|---|
| Milli-Q water ($dH_2O$) | 410 mL | | | |
| 1 M Potassium phosphate buffer, pH 7.2 | 50 mL | 10% (v/v) | | |
| TYG salts solution | 20 mL | 4% (v/v) | | |
| Tween 80 | 1 mL of 25% (v/v) | 0.05% (v/v) | | |
| SCFA supplement | 1.4 mL | 0.28% (v/v) | | |
| 0.8% (w/v) $CaCl_2$ | 500 µL | | | |
| $FeSO_4 \cdot 7H_2O$ | 500 µL of 0.4 mg/mL | | | |
| Resazurin | 2 mL of 0.25 mg/mL | 0.000001% (w/v) | Sigma | Sigma (R2127) |
| Trypticase Peptone (BBL) | 5 g | 1% (w/v) | BD | BD (211921) |
| Yeast Extract (Bacto) | 2.5 g | 0.5% (w/v) | BD | BD (212750) |
| Meat extract | 2.5 g | 0.5% (w/v) | Sigma | Sigma (70164) |
| L-Cysteine hydrochloride | 0.25 g | 0.05% (w/v) | Sigma | Sigma (C1276) |

| | | | | |
|---|---|---|---|---|
| D-(+)-Glucose | 1 g | 0.2% (w/v) | Sigma | Sigma (G8270) |
| D-(+)-Cellobiose | 0.5 g | 0.1% (w/v) | Sigma | Sigma (C7252) |
| D-(+)-Maltose monohydrate | 0.5 g | 0.1% (w/v) | Sigma | Sigma (M5885) |
| D-(-)-Fructose | 0.5 g | 0.1% (w/v) | Sigma | Sigma (F0127) |
| Soluble starch | 0.25g | 0.05% (w/v) | | |

**Table 2**

| Component | Amount(in 500 mL) | Final conc. | Mfr. | Vendor (cat. #) |
|---|---|---|---|---|
| Vitamin K solution | 500 µL of 1 mg/mL | 0.000001% (w/v) | Sigma | Sigma (M5625) |
| Trace Mineral Supplement | 5 mL | 1% (v/v) | ATCC | ATCC (MD-TMS) |
| Vitamin Supplement | 5 mL | 1% (v/v) | ATCC | ATCC (MD-VS) |
| Histidine-Hematin | 500 µL | | | |

### STOCK SOLUTION RECIPES

#### 1 M potassium phosphate buffer, pH 7.2

1. Prepare 1 M $KH_2PO_4$ (monobasic).
   68.045 g $KH_2PO_4$ (anhydrous, f.w. = 136.09) in Milli-Q water to 500 mL
2. Prepare 1 M $K_2HPO_4$ (dibasic).
   174.18 g $K_2HPO_4$ (anhydrous, f.w. = 174.18) in Milli-Q water to 1 L
3. Add monobasic to dibasic to achieve pH 7.2.

(You typically need ~430 mL monobasic added to 1 L dibasic)

*Tip: When preparing this solution, add the potassium phosphate powders to an actively stirring volume of water. Attempting to add water to a large mass of powder will result in the formation of a difficult to dissolve clump of material at the bottom of the bottle.*

#### Vitamin K solution

Dissolve 40 mg menadione (Vitamin $K_3$, Sigma M5625) in 40 mL 100% EtOH.

#### TYG salts solution

$MgSO_4 \cdot 7H_2O$ (Sigma 230391) 0.5 g
$NaHCO_3$ (Sigma S5761) 10.0 g
NaCl (Sigma S7653) 2.0 g

| | |
|---|---|
| Milli-Q water | to 1 L |

**$FeSO_4 \cdot 7H_2O$ (0.4 mg/mL)**

Dissolve 40 mg $FeSO_4 \cdot 7H_2O$ (Sigma F8633) in 100 mL Milli-Q water.

**0.8% (w/v) $CaCl_2$**

Dissolve 0.4 g $CaCl_2 \cdot 2H_2O$ (Sigma C7902) in 50 mL Milli-Q water.

**Resazurin anaerobic indicator (0.25 mg/mL)**

1. Dissolve 25 mg resazurin (Sigma R2127) in 100 mL distilled $H_2O$.
2. Store protected from light at 4 °C.

**Histidine-Hematin**

1. Prepare 0.2 M histidine, pH 8.0

    · Mix 4.2 g Histidine-HCl monohydrate (Sigma H7875) in 80 mL Milli-Q water.
    · Adjust the pH from 4 to 8 with 10 N NaOH (the histidine will go into solution as the pH rises).
    · Bring the final volume to 100 mL with Milli-Q water.

2. Mix 12 mg hematin (Sigma H3281) with 10 mL of 0.2 M histidine, pH 8.0. Dissolve by end-over-end rotation or vigorous shaking for several hours. Filter-sterilize using 0.2 µm filter.

**SCFA supplement**

| | |
|---|---|
| Acetic acid, glacial (Sigma A6283) | 17 mL |
| Propionic acid (Sigma P5561) | 6 mL |
| Butyric acid (Sigma B103500) | 4 mL |
| Isovaleric acid (Sigma 129542) | 1 mL |

**Salyer's Minimal Medium (SMM) preparation protocol**

This recipe was adapted from multiple references[8,9] that give slightly different concentrations for various components. The media was prepared and used as described in the supplemental methods.

Mineral salts (Table 3 below) and hematin solution can be prepared, filter sterilized, and stored at 4°C for at least 12 months. Carbon sources and nitrogen sources (critically if using glutamine) should be prepared the day of use.

Steps:

1. Prepare mineral salts, hematin, vitamin K3 solution, iron sulfate solution, and vitamin $B_{12}$ solution.
2. Prepare SMM base according to Table 1 below. Filter sterilize and place into anaerobic chamber for reduction. May take 48 hours given paucity of reductants.
3. Prepare carbon, nitrogen, and sulfur sources immediately prior to use, filter sterilize, and add to reduced SMM base according to Table 2 below.

**Table 1 – SMM Base**

| Component | Amount (in 100 mL) |
|---:|---:|
| $dH_2O$ | 60.84 ml |
| 1 M KPO4, pH 7.2 | 10 ml |
| Vitamin K3 solution, 1mg/ml | 0.01 ml |
| FeSO4.7H2O, 0.4 mg/ml | 1 ml |
| Vitamin $B_{12}$, 0.01 mg/ml | 0.05 ml |
| $Na_2CO_3$ | 0.1 g |

**Table 2 – SMM Complete**

| Component | Amount (in 100 mL) |
|---:|---:|
| SMM base | 71.9 ml |
| Carbon source (50x) | 2 ml |
| Mineral salts | 5 ml |
| Reduced sulfur source (100X) | 1 ml |
| Hematin solution | 0.1 ml |
| Nitrogen source (5X) | 20 ml |

**Table 3 – Mineral salts**

| Component | Amount (in 1L) |
|---|---|
| NaCl | 18 g |
| $CaCl_2.2H_2O$ | 0.53 g |
| $MgCl_2.6H_2O$ | 0.40 g |
| $MnCl_2.4H_2O$ | 0.20 g |
| $CoCl_2.6H_2O$ | 0.20 g |
| $dH_2O$ | 1 L |

Carbon source at a final concentration of 27.7 mM (50X=1.385M). Here we used Glucose

| Component | Amount (in 50mL) |
|---|---|
| $dH_2O$ | 50 ml |
| Glucose | 12.48 g |

Hematin at a 100X concentration. To prepare this, first mix 0.4g NaOH in 100ml $dH_2O$, then add 0.5g Hemin or Hematin.

| Component | Amount (in 100ml) |
|---|---|
| 0.1 M NaOH<br>(0.4g NaOH in 100ml $dH_2O$) | 100 ml |
| Hemin | 0.5 g |

Reduced sulfur source at a final concentration of 4.12 mM (100X = 412.7mM). Here we used both cysteine and sodium sulfide.

| Component | Amount (in 20mL) |
|---|---|
| $dH_2O$ | 20 ml |
| Cysteine | 1 g |
| $dH_2O$ | 20 ml |
| Na2S | 0.644 g |

Nitrogen sources at a final concentration of 10mM (5X = 50 mM). Ammonium sulfate was assumed to produce two equivalents and all others one equivalent of $NH_4$.

| **Component** | **Amount (in 30mL)** |
|---:|---:|
| $dH_2O$ | 30 ml |
| $(NH_4)_2SO_4$ | 0.1 g |
| $dH_2O$ | 30 ml |
| Glutamate | 0.22 g |
| $dH_2O$ | 30 ml |
| Glutamine | 0.22 g |
| $dH_2O$ | 30 ml |
| Asparagine | 0.2 g |
